# DYT-TOR1A Subcellular Proteomics Reveals Selective Vulnerability of the Nuclear Proteome to Cell Stress

**DOI:** 10.1101/2021.07.10.451533

**Authors:** Kunal Shroff, Zachary F. Caffall, Nicole Calakos

**Affiliations:** Department of Neurology, Duke University Medical Center; Department of Neurobiology, Duke University Medical Center; Department of Cell Biology, Duke University Medical Center; Department of Duke University Medical Center, Duke Institute for Brain Sciences, Duke University Durham, North Carolina, USA

**Keywords:** Dystonia, Subcellular fractionation, TorsinA, Compartment-specific proteome, Stress Response, Movement disorder

## Abstract

TorsinA is a AAA^+^ ATPase that shuttles between the ER lumen and outer nuclear envelope in an ATP-dependent manner and is functionally implicated in nucleocytoplasmic transport. We hypothesized that the DYT-TOR1A dystonia disease-causing variant, ΔE TorsinA, may therefore disrupt the normal subcellular distribution of proteins between the nuclear and cytosolic compartments. To test this hypothesis, we performed proteomic analysis on nuclear and cytosolic subcellular fractions from DYT-TOR1A and wildtype mouse embryonic fibroblasts (MEFs). We further examined the compartmental proteomes following exposure to thapsigargin (Tg), an endoplasmic reticulum (ER) stressor, because DYT-TOR1A dystonia models have previously shown abnormalities in cellular stress responses. Across both subcellular compartments, proteomes of DYT-TOR1A cells showed basal state disruptions consistent with an activated stress response, and in response to thapsigargin, a blunted stress response. However, the DYT-TOR1A nuclear proteome under Tg cell stress showed the most pronounced and disproportionate degree of protein disruptions – 3-fold greater than all other conditions. The affected proteins extended beyond those typically associated with stress responses, including enrichments for processes critical for neuronal synaptic function. These findings highlight the advantage of subcellular proteomics to reveal events that localize to discrete subcellular compartments and refine thinking about the mechanisms and significance of cell stress in DYT-TOR1A pathogenesis.

**Highlights:**

- The DYT-TOR1A nuclear proteome under cell stress showed 3-fold greater protein disruptions.
- DYT-TOR1A MEFs show basal proteome alterations consistent with cell stress.
- Thapsigargin modulation of WT stress-responsive proteins is blunted in DYT-TOR1A MEFs.
- TorsinB was identified as part of the cell-stress responsive proteome in WT MEFs.

**Graphical Abstract:** 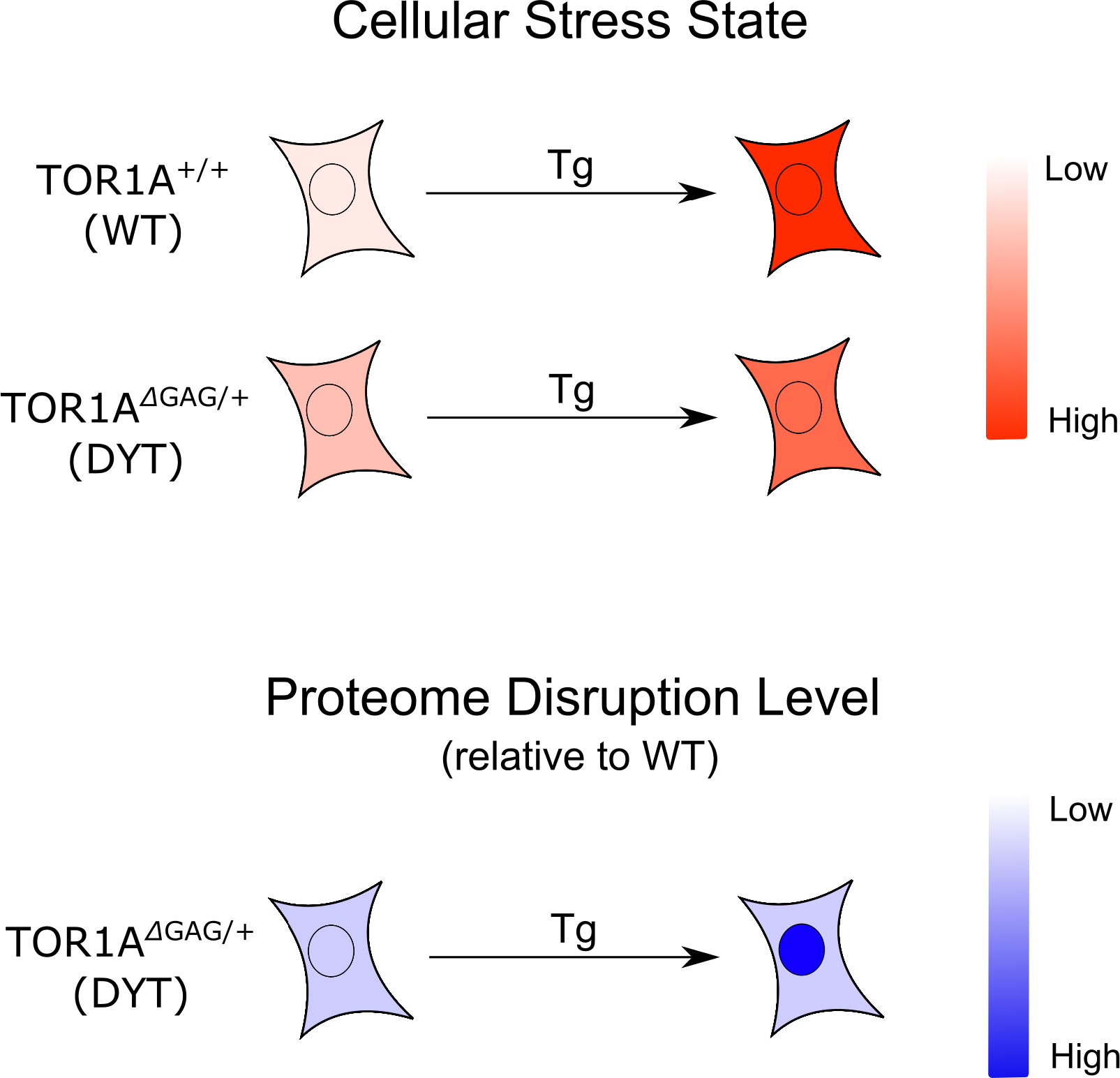

## Introduction

Dystonia is a neurological movement disorder characterized by involuntary twisting and abnormal postures of the limbs, trunk, and/or face (Tarsy & Simon, 2006). Causes for dystonia are diverse, ranging from exposure to anti-psychotic medications to neurodegenerative diseases (Balint et al., 2018; Bressman, 2004; Jankovic & Tintner, 2001; van Harten et al., 1999). DYT-TOR1A dystonia is a rare inherited autosomal dominant form of the disorder that is caused by an in-frame trinucleotide deletion in the *Tor1a* coding sequence (n. ΔGAG, p. ΔE) and leads to an early-onset, generalized dystonia (Ozelius et al., 1997).

Since the discovery of *Tor1a* as the causal gene for DYT-TOR1A dystonia, a number of groups have characterized the function of the encoded protein, TorsinA, in its normal and mutant forms. TorsinA is a member of the AAA+ ATPase family of proteins. Although typically TorsinA is predominantly in the lumen of the endoplasmic reticulum, perturbations preventing ATP hydrolysis result in a prominent outer nuclear envelope distribution (Naismith et al., 2004). These observations led to a model in which the protein shuttles between these two compartments in an ATP hydrolysis-dependent manner (Naismith et al., 2004). Later studies identified two TorsinA binding partners, LAP1 and LULL1, which localize to the nuclear envelope and ER membrane, respectively (Chalfant et al., 2019; Chase et al., 2017; Esra Demircioglu et al., 2016; Goodchild & Dauer, 2005; M. T. Jungwirth et al., 2011; Laudermilch et al., 2016; Nery et al., 2008; Saunders et al., 2017; Vander Heyden et al., 2009). Thus, it was hypothesized that TorsinA likely played a role in each of these compartments. More recent studies implicate TorsinA in regulation of nucleocytoplasmic transport (Chalfant et al., 2019; Ding et al., 2021; György et al., 2018; Jokhi et al., 2013; Laudermilch et al., 2016; Rampello et al., 2019; VanGompel et al., 2015). Defects in nucleoporin localization and nuclear import kinetics have been described in association with mutations in OOC-5, a TorsinA homolog in *C. elegans,* and in *Tor1a* in mammalian neuronal cultures and knockout mouse models (Chalfant et al., 2019; Pappas et al., 2018; VanGompel et al., 2015). In addition, impaired nuclear egress functions were found in both DYT-TOR1A patient fibroblasts and cultured DYT-TOR1A mouse neurons (György et al., 2018). These defects may be related to TorsinA’s role in regulating nuclear budding, a process essential for nuclear egress of mega-ribonucleoproteins (megaRNPs) (Jokhi et al., 2013; Speese et al., 2012). Collectively, these findings suggest that TorsinA plays a role in regulating nucleocytoplasmic transport, and that the DYT-TOR1A-associated TorsinA mutation, ΔE, impairs these functions.

In addition to nucleocytoplasmic transport defects, our lab and several others have found that multiple DYT-TOR1A dystonia model systems exhibit altered cellular stress response pathways (Beauvais et al., 2016, 2018; Chen et al., 2010; Cho et al., 2014; Nery et al., 2011; Rittiner et al., 2016; Zacchi et al., 2014; Zhao et al., 2016). It is currently unknown how the DYT-TOR1A mutation specifically causes these cellular stress response defects. However, alleviation of these defects by pharmacological and genetic approaches has been shown to improve DYT-TOR1A associated cellular phenotypes (Rittiner et al., 2016), suggesting that cellular stress responses may play an important role in DYT-TOR1A disease pathophysiology.

Proteomics approaches have been previously used to study the consequences of the DYT-TOR1A causative mutation in whole cell and tissue lysate preparations (Beauvais et al., 2016, 2018; Martin et al., 2009). These approaches have successfully identified disease model-associated protein defects. However, because many subcellular compartments such as the nucleus comprise only a proportionally small part of the total proteome, compartment-specific defects are likely to be overlooked by such approaches. Recent studies of amyotrophic lateral sclerosis demonstrate the potential for nucleocytoplasmic proteomic analyses by identifying nucleus-specific defects in RNA transport and cytosol-specific defects in protein translation and folding (J. E. Kim et al., 2017; Ortega et al., 2020). In the present study, we adopt a similar nucleocytoplasmic fractionation technique alongside quantitative proteomics to address whether the DYT-TOR1A mutation causes subcellular compartment-specific proteomic disruptions.

Our results identify compartment- and stress-specific disruptions associated with the DYT-TOR1A genotype that include disruptions of proteins that are normally cell stress modulated and of TorsinA and TorsinB levels and localization. The most striking result however was that, despite our leading hypothesis that defects in nucleocytoplasmic transport might affect both compartments, the DYT-TOR1A mutation was found to cause the most pronounced and disproportionate insult to the nuclear proteome and selectively under cell stress. This result indicates that ΔE TorsinA causes a particular vulnerability to the integrity of the nuclear proteome in the face of cellular stressors.

## Materials and Methods

### Animals

DYT-TOR1A knock-in mice *(Tor1a*^ΔGAG/+^)(courtesy of Dr. W. Dauer, University of Michigan) (Goodchild et al., 2005) on C57BL/6 background were bred in standard housing conditions with food and water provided ad libitum. All procedures were approved by the Duke University Institutional Animal Care and Use Committee (IACUC).

### Mouse Embryonic Fibroblast (MEF) Extraction, Isolation, and Immortalization

To produce DYT-TOR1A model MEF and WT control MEF cell lines, female wildtype C57BL/6 mice were crossed with male heterozygous DYT-TOR1A knock-in mice (*Tor1a*^ΔGAG/+^) on a C57BL/6 background. MEF extraction occurred with minor modifications from the protocol as described in (Jozefczuk et al., 2012). Three DYT-TOR1A MEF lines and three WT MEF lines were produced from littermates of a single litter. The pregnant dam was euthanized at approximately 14 days post-coitum using isoflurane followed by decapitation. The uterine horns were dissected out and rinsed in 70% (v/v) ethanol and PBS (Gibco, Invitrogen) before placing into a Petri dish containing PBS (Gibco, Invitrogen). Each individual embryonic sac was separated from the uterine horns and placenta, and then placed into a separate dish containing PBS. Each embryo was dissected out of the embryonic sac and its head and red organs were removed. The remaining embryonic tissue was placed into a clean Petri dish where it was minced with a sterile razor blade. 1 mL of 0.05% trypsin/EDTA (Gibco, Invitrogen) was added to each dish. The mixture was transferred into a 15 mL Falcon tube and incubated at 37 °C for 30 minutes. After each 10 minutes of incubation, MEFs were dissociated via pipetting. Trypsin was inactivated by adding 2 mL of fetal bovine serum-containing media (“MEF media” described in Cell Culture section below) to each tube. The MEFs were then centrifuged at 500 x g for 5 minutes. The supernatant was removed, and the cell pellet was resuspended in 10 mL of warm MEF media. This solution was then plated on TC dishes coated in 1% Matrigel (Corning). After 2 passages, the MEFs were genotyped and subsequently immortalized via the SV40 T antigen as described in (H. Harding, 2003). Cell lines were used within 5 passages.

### Genotyping

All genotyping was conducted as previously described in (Goodchild et al., 2005).

### Cell Culture

MEFs were grown in MEF media which consisted of 500 mL of DMEM, high glucose, pyruvate (Thermo Fisher, #11995), 50 mL of Fetal Bovine Serum, 5 mL of Antibiotic-Antimycotic (Gibco, Invitrogen), 5 mL of 200 mM L-Glutamine (Gibco, Invitrogen), 5 mL of MEM Non-Essential Amino Acids Solution (Gibco, Invitrogen), and 500 μL of 2-Mercaptoethanol (Sigma-Aldrich). MEFs were grown in incubators at 37 °C/5% CO2.

### Thapsigargin Treatment and Subcellular Fractionation

Three separate experiments were performed exposing MEFs to the cell stressor, thapsigargin (Tg). During each experiment, a pair of MEF lines (1 WT and 1 DYT-TOR1A) was treated with either 1 μM Tg dissolved in DMSO or an equivalent volume of DMSO (Vehicle control, Veh). After six hours of treatment at 37 °C, the MEFs were subcellularly fractionated into nuclear and cytosolic fractions. MEFs for subcellular fractionation were acquired through trypsinization from 90% confluent TC plates. Subcellular fractionation of MEFs was then carried out as described in (Suzuki et al., 2010). Briefly, the procedure involves a weak and brief detergent extraction (0.01% NP40, 3 min., room temperature), centrifugation to collect the supernatant (cytosolic fraction) and then further solubilization in the same buffer alongside sonication to penetrate the double bilayer membranous nuclear compartment, with a second centrifugation and supernatant collection for the nuclear fraction.

### Western Blotting

Lysates for Western analysis were produced either through the subcellular fractionation protocol described earlier or via a whole-cell lysate produced with RIPA buffer-induced cell lysis. Protein content from each lysate was determined via Bicinchoninic Acid (BCA) assay. Samples were prepared such that each sample contained an equal mass of protein and 1x Laemmli buffer. Samples were reduced and denatured with 2-mercaptoethanol and incubated at 97 °C for 5 minutes. Equal volumes of sample were loaded into the wells of an SDS-PAGE gel along with a protein ladder. After 45 minutes electrophoresis at 175 V, the protein within the gel was transferred to a nitrocellulose membrane.

Following transfer, the membrane was blocked in 5% BSA solution prepared in TBST for 1 hour at room temperature. The membrane was then incubated overnight on a shaker at 4 °C in blocking solution amended with the primary antibody [anti-Lamin B1 (Abcam; ab16048; 1:1000), anti-GAPDH (Abcam; ab9485; 1:1000), anti-Na+/K+ - ATPase (Santa Cruz Biotechnology; sc-21712; 1:500), anti-BiP (Santa Cruz Biotechnology; sc-13968; 1:500)]. Following primary incubation, the membrane was washed three times with TBST for 5-10 minutes each time. The membrane was then re-blocked in blocking solution for 1 hour. Membranes were then placed in blocking solution with secondary antibody [Alexa Fluor 790 Goat anti-Rabbit and/or Alexa Fluor 680 Goat anti-Mouse (Thermo Fisher)] at a dilution of 1:1000. The membrane was incubated in the secondary solution for 1 hour before being washed three times in TBST for 5-10 minutes per wash. The membrane was imaged on a LI-COR Odyssey Imaging System. The visualized bands were quantified using ImageJ.

### Immunofluorescent Staining

MEF lines (three WT and three DYT-TOR1A) were plated into individual wells on a 96-well plate, such that each line was plated into eight wells. Four of the wells for every line were treated with Veh and the other four wells for each line were treated with 1 μM Tg for six hours. Following treatment, the wells were fixed with 4% paraformaldehyde, permeabilized and blocked with blocking solution (0.1% Triton-X 100 in PBS, 1% bovine serum albumin, 10% normal donkey serum) for 1 hour. The wells were then stained with a primary antibody [anti-Tbce (Thermo-Fisher, PA5-100346, 1:200), anti-Pds5B (Thermo-fisher; PA5-59029; 1:500)] overnight at 4°C. Following three washes with wash buffer solution (0.1% BSA in PBS), secondary staining was conducted using Hoechst 33342 (MilliporeSigma; 1:1000) to stain the nucleus and donkey Anti-rabbit Alexa Fluor 488 (Life Technologies; 1:1000) for 1 hour at room temperature. Following three additional washes with wash buffer solution, the wells were filled with dilution buffer (1% BSA, 1% normal donkey serum, 0.3% Triton X-100, and 0.01% sodium azide in PBS). Sixteen imaging fields from each well were acquired at a magnification of 20x from a Thermofisher CX5 HC imager. Images were acquired in both the blue and green channel to identify the cell nuclei and quantify the protein of interest. Following image acquisition, images were analyzed by CellProfiler 3.1.9 (McQuin et al., 2018).

### Immunofluorescence Quantification and Data Analysis

Nuclei were identified via the blue channel Hoechst stain. The cytosol was identified via the green channel protein immunofluorescence stain and nuclear position information as determined from the blue channel Hoechst stain. Nuclear fluorescence was quantified by integrating the intensity of the green channel protein immunofluorescence across pixels identified as being part of the nucleus. Cytosolic fluorescence was quantified by integrating the intensity of the green channel protein immunofluorescence across each pixel within the region identified as being part of the cytosol.

Puncta were identified via a modified speckle counting pipeline developed by CellProfiler (McQuin et al., 2018). Analyses were conducted on image masks containing only the nuclei, as well as image masks containing only the cytosol to quantify puncta in each of the subcellular compartments. Puncta frequency was determined by taking the number of speckles identified per image and then dividing by the number of nuclei or cytosolic areas within that image. Puncta intensity was determined by integrating the intensity of all the puncta within the field and dividing by the number of nuclei or cytosolic areas within that image.

Each quantified value (nuclear fluorescence, cytosolic fluorescence, puncta frequency, puncta intensity) was calculated for each of the sixteen fields imaged per well and these sixteen values were averaged to produce a single mean value for each well. Unpaired t-tests were used to compare mean values from the four biological replicate wells across both genotypes with and without stress treatment.

### Quantitative LC/MS/MS and Proteomic Analysis

Twenty-four samples in total were submitted to the Duke Proteomics and Metabolomics Shared Resource (two subcellular fractions from each of the six MEF lines treated with either Tg or Veh). While the fractions were collected over three separate cell culture experiments, they were all analyzed within a single liquid chromatography with tandem mass spectroscopy (LC/MS/MS) experiment. Fractions were first normalized to 10 μg and spiked with undigested casein at a total of 20, 30, or 40 fmol/μg, then reduced with 10 mM dithiothreitol for 30 min at 80 °C, and alkylated with 20 mM iodoacetamide for 30 min at room temperature. Next, they were supplemented with a final concentration of 1.2% phosphoric acid and 741 μL of S-Trap (Protifi) binding buffer (90% MeOH/100mM TEAB). Proteins were trapped on the S-Trap, digested using 20 ng/μL sequencing grade trypsin (Promega) for 1 hour at 47°C, and eluted using 50 mM TEAB, followed by 0.2% FA, and lastly using 50% ACN/0.2% FA. All fractions were then lyophilized to dryness and resuspended in 20 μL 1% TFA/2% acetonitrile containing 12.5 fmol/μL yeast alcohol dehydrogenase (ADH_YEAST). Three QC Pools were created: 1) 3 μL from each of the nuclear fractions, 2) 3 uL from each of the cytosolic fractions 3) 3 μL from each of all of the fractions, both nuclear and cytosolic. All QC Pools were run periodically randomly interspersed throughout the test fractions.

Quantitative LC/MS/MS was performed on 2 μL of each fraction, using a nanoAcquity UPLC system (Waters Corp.) coupled to a Thermo Orbitrap Fusion Lumos high resolution accurate mass tandem mass spectrometer (Thermo) via a nanoelectrospray ionization source. Briefly, the fraction was first trapped on a Symmetry C18 20 mm × 180 μm trapping column (5 μL/min at 99.9/0.1 v/v water/acetonitrile), after which the analytical separation was performed using a 1.8 μm Acquity HSS T3 C18 75 μm × 250 mm column (Waters Corp.) with a 90-min linear gradient of 5 to 30% acetonitrile with 0.1% formic acid at a flow rate of 400 nanoliters/minute (nL/min) with a column temperature of 55 °C. Data collection on the Fusion Lumos mass spectrometer was performed in a data-dependent acquisition (DDA) mode of acquisition with a r=120,000 (@ m/z 200) full MS scan from m/z 375 – 1500 with a target AGC value of 2e5 ions. MS/MS scans were acquired at Rapid scan rate (Ion Trap) with an AGC target of 5e3 ions and a max injection time of 100 milliseconds. The total cycle time for MS and MS/MS scans was 2 seconds. A 20s dynamic exclusion was employed to increase depth of coverage. The total analysis cycle time for each fraction injection was approximately 2 hours.

Following 35 total UPLC-MS/MS analyses (excluding conditioning runs, but including 3 replicate QC Pool, 4 replicate nuclear and 4 replicate cytosolic Pool injections), data was imported into Proteome Discoverer 2.2 (Thermo Scientific Inc.), and analyses were aligned based on the accurate mass and retention time of detected ions (“features”) using Minora Feature Detector algorithm in Proteome Discoverer. Protein levels are reported in arbitrary units (a.u.) based on the relative peptide abundance measures which were calculated by area-under-the-curve (AUC) of the selected ion chromatograms of the aligned features across all runs. The MS/MS data was searched against the SwissProt *M. musculus* database (downloaded in Apr 2017) and an equal number of reversed-sequence “decoys” for false discovery rate determination. Mascot Distiller and Mascot Server (v 2.5, Matrix Sciences) were utilized to produce fragment ion spectra and to perform the database searches. Database search parameters included fixed modification on Cys (carbamidomethyl) and variable modifications on Meth (oxidation) and Asn and Gln (deamidation). Peptide Validator and Protein FDR Validator nodes in Proteome Discoverer were used to annotate the data at a maximum 1% protein false discovery rate.

Missing values were imputed after sample loading normalization in the following manner. If less than half of the values are missing across all samples, values are imputed with an intensity derived from a normal distribution defined by measured values within the same intensity range (20 bins). If greater than half values are missing for a peptide across all samples and a peptide intensity is > 5e6, then it was concluded that peptide was misaligned and its measured intensity is set to 0. All remaining missing values are imputed with the lowest 5% of all detected values. All analyses presented here are based on these normalized values. The complete proteomic dataset has been deposited with Mendeley data and is further detailed in a Data In Brief accompanying article.

### Data Analysis and Statistical Analysis

Throughout the data collection phase of the study, cell genotype and stress-treatment conditions were blinded variables. Unblinding occurred upon return of processed proteomic data.

For proteomic data analysis, proteins represented by only a single peptide were removed from the data set prior to further analysis to reduce the number of Type 1 errors. Unpaired t-test p-values and fold changes for each protein were calculated for each comparison. P-values and fold changes were calculated using GSEA software from the Broad Institute via a Student’s t-test (Mootha et al., 2003; Subramanian et al., 2005). Proteins that showed an uncorrected p-value less than 0.05 and a DYT-TOR1A/WT or WT/DYT-TOR1A ratio greater than 1.5 (DYT-TOR1A/WT fold change greater than ±log_2_(0.585)) were considered as the “top hits” for further analysis. Metascape was used to conduct a Gene Ontology analysis on the top hits (Zhou et al., 2019). Top hits were analyzed using the entire discovered proteome as the background to consider for enrichment.

All other statistical testing used unpaired t-tests calculated by GraphPad Prism version 8.3.1 for MacOS unless otherwise indicated.

## Results

### Subcellular Fractionation of Mouse Embryonic Fibroblasts (MEFs)

Immortalized murine embryonic fibroblast cell lines were prepared from *Tor1a*^ΔGAG/+^ (genotype hereafter abbreviated as DYT-TOR1A or DYT) and wildtype (WT) littermate embryos according to standard methodology (Methods). Cultures from 3 independent lines for each genotype were grown to confluence and then treated with either 1 μM thapsigargin (Tg) or vehicle (Veh) for six hours prior to harvesting (Fig. 1A). Nuclear and cytosolic cellular fractions were prepared according to previously described methods based on serial exposure to a mild detergent extraction that does not significantly solubilize nuclear membranes, but is sufficient to penetrate plasma membrane and enable extraction of cytosolic components and organelles, followed by a nuclear membrane-solubilizing sonication step (Suzuki et al., 2010). Western analysis was performed to confirm that markers for the nuclear (Lamin B1) and the cytosolic (GAPDH, BiP, Na,K-ATPase α1) subcellular compartments were differentially distributed in the fractions as predicted in both genotypes (Fig. 1B and Fig. S1). Nuclear and cytosolic fractions prepared from 3 independent cell lines for each genotype were then analyzed by quantitative liquid chromatography-tandem mass spectrometry (LC/MS/MS).

**Fig. 1.**
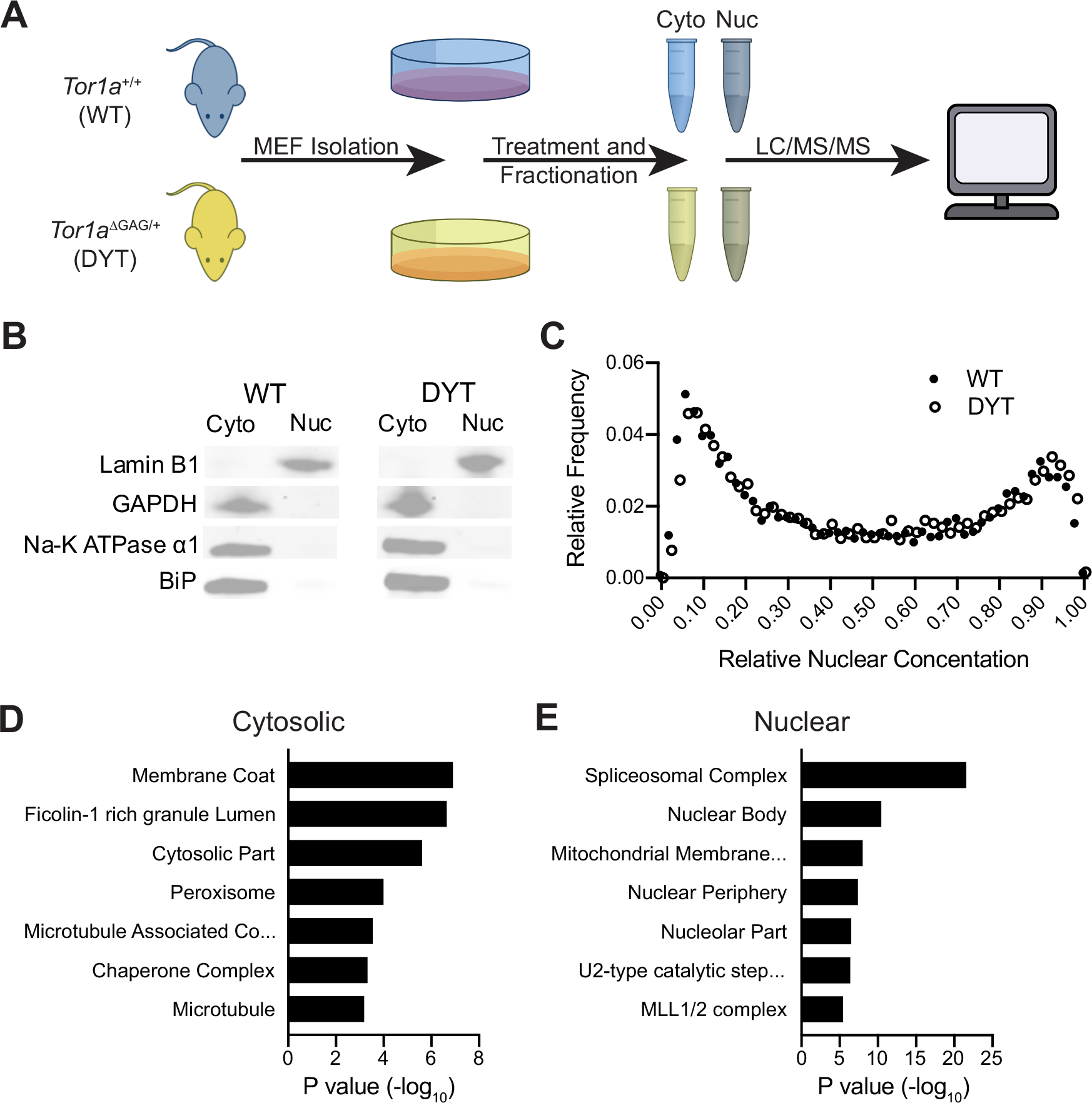
Subcellular fractionation enriches for cytosolic and nuclear components similarly in both WT and DYT-TOR1A cell lines. (A) Experimental design schematic. Mouse embryonic fibroblasts (MEFs) were isolated from DYT-TOR1A heterozygous knock-in (*TOR1A*^ΔGAG/+^) mice and wildtype litter mates. MEF lysates were fractionated into cytosolic (Cyto) and nuclear (Nuc) fractions and then subjected to quantitative differential proteomics analysis. (B) Representative Western blots of subcellular fraction markers in WT and DYT-TOR1A cell lines. The nuclear membrane marker, Lamin B1, is predominately sequestered in the nuclear fraction while GAPDH, Na-K ATPase α1, and BiP (cytosolic, plasma membrane, and endoplasmic reticulum resident proteins, respectively) are enriched in the cytosolic fraction. (C) Frequency distribution of calculated relative nuclear concentration (RNC) values across the entire proteome from WT (solid circles) and DYT-TOR1A (open circles) samples. RNC values are calculated for each individual protein by taking the protein abundance within the nuclear fraction and dividing by the sum of the nuclear and cytosolic fraction protein abundances (adapted from Wühr et al., 2015). (D-E) Gene Ontology analysis of proteins enriched within the nuclear (D) and cytosolic fraction (dataset of combined genotypes using threshold of p<0.001 by t-test).

A total of 4801 proteins were detected across all samples. Of those, 3921 proteins had at least two distinct peptides and mapped to a unique mouse gene identifier using GSEA software; these proteins were used for subsequent analyses. Over 90% of proteins were identified in both subcellular fractions and treatment conditions (Fig. S2A). Recognizing that most proteins are present in both subcellular compartments to varying degrees, we next calculated the relative nuclear concentration (RNC) (nuclear level/total_nuclear + cytosolic_) for each of the 3921 proteins. The RNC has been used previously to characterize the proteome and demonstrated only a small fraction of proteins being almost exclusive to nuclear or cytosolic fractions, with a majority of proteins having intermediate RNC values (Wühr et al., 2015). The nucleocytoplasmic distribution of proteins in our samples is consistent with those prior observations and was similar across the two genotypes (Fig. 1C). In addition, we performed standard bioinformatic analysis using Gene Ontology (GO) on the nuclear and cytosolic fraction proteomic datasets to determine whether enrichments characteristic of nuclear and cytosolic components were detected in the corresponding fractions (For ease of presentation, results of both genotypes were combined. Individual genotype analyses yielded similar conclusions, data not shown). The top GO terms associated with proteomics of the cytosolic fractions included “cytosolic part” (Fig. 1D). In addition to cytosolic proteins, we also observed enrichment for cytosolic vesicle membrane proteins as shown by the strong enrichment of the GO terms “membrane coat” and “Ficolin-1 rich granule lumen”. Conversely, the top GO terms associated with nuclear fractions included “nuclear body” (Fig. 1E). We further noted that GO analysis of the nuclear fraction also included “mitochondrial membrane part” suggesting that mitochondria may be preferentially extracted with the nuclear fraction. Together, these characterizations establish that components of the nucleus and cytosol are relatively enriched in the nuclear and cytosolic fractions, respectively.

### DYT-TOR1A Genotype-Dependent Subcellular Proteome Differences

To identify proteome differences caused by the DYT-TOR1A genotype, an average fold change (FC) and p-value were calculated for each protein by comparing levels between WT and DYT-TOR1A samples under basal conditions (Veh) (n = 3 independent biological replicates per genotype). Using thresholds of an uncorrected p-value of less than 0.05 and fold change of ±1.5 FC, which corresponds to a 50% increase in DYT-TOR1A levels relative to WT or WT levels relative to DYT-TOR1A, we identified 152 proteins with genotype-dependent differences in the cytosolic fractions (Fig. 2A). A similar number of differences were identified in the nuclear fractions (Fig. 2B). There was less than 3% overlap between the differentially affected proteins in the nuclear and cytosolic samples (Fig. S2B). Gene Ontology analysis of differentially affected proteins revealed enrichment of proteins associated with mitochondrion organization and ATP metabolism (Fig. S3). This GO term enrichment was present in both the nuclear and cytosolic DYT-TOR1A-disrupted protein datasets (Fig. S3). These results are consistent with processes that have been previously implicated in DYT-TOR1A dystonia (Beauvais et al., 2016; Martin et al., 2009).

**Fig. 2.**
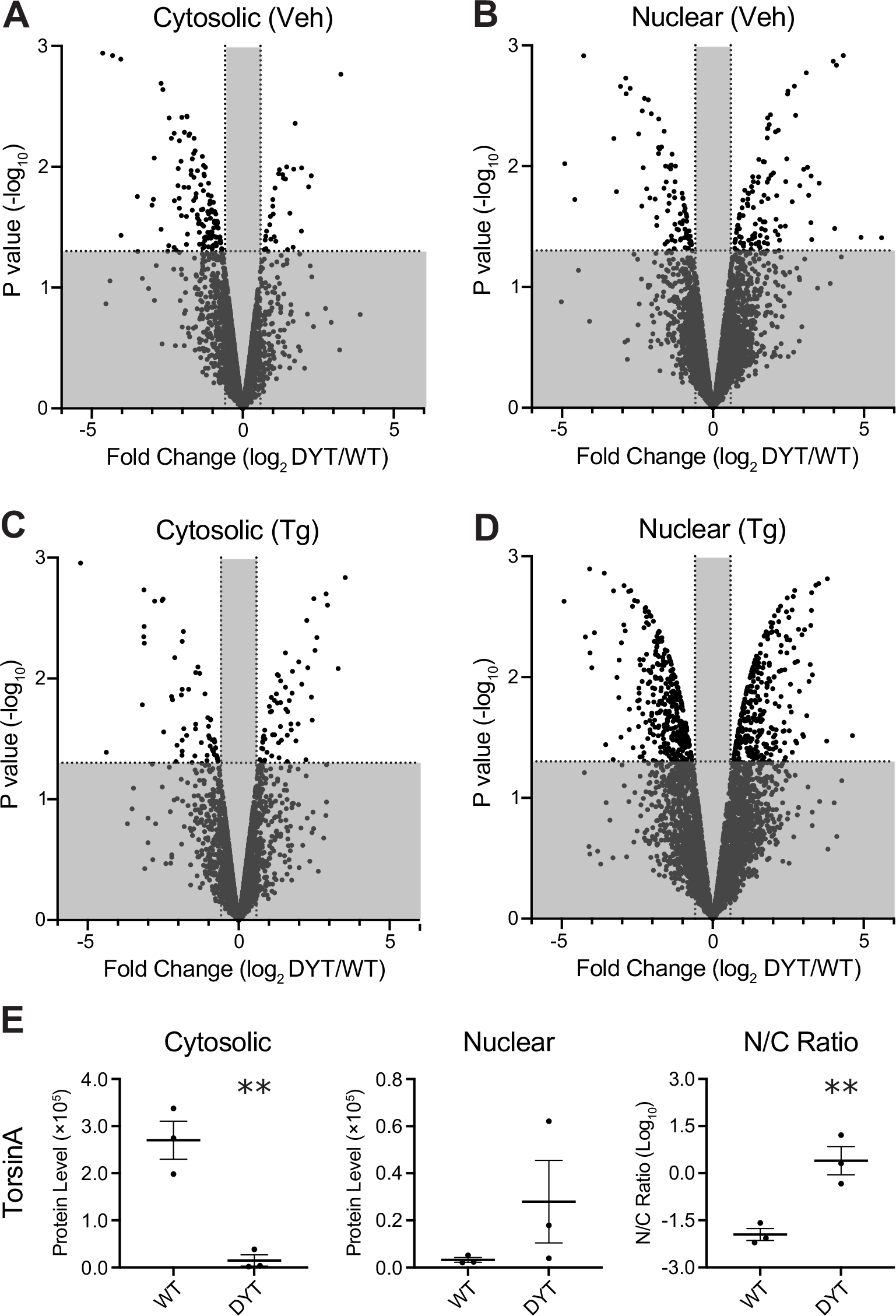
Compartment-specific proteome disruptions associated with DYT-TOR1A MEFs. (A-B) Volcano plots showing proteome-wide differences in protein abundances between DYT-TOR1A and WT cytosolic (A) and nuclear (B) fractions under basal conditions (Veh). Differences in protein abundance are represented as fold change (using log2 transform) and p-value is calculated by unpaired t-test for each protein (n=3 biological replicates). Horizontal dashed line indicates p-value less than 0.05. Vertical dashed lines indicate fold changes of ±1.5. (C-D) Corresponding volcano plots for fractions from thapsigargin-treated (Tg) cells. (E) Relative TorsinA peptide levels (a.u.) in cytosolic (Veh) and nuclear (Veh) fractions and the nuclear:cytosolic (N/C) ratio. Error bars indicate S.E.M.. For all comparisons, n = 3 biological replicates, p-value determined by unpaired t-test (**p<0.01).

Prior studies have also described numerous cellular stress response defects in DYT-TOR1A dystonia models (Beauvais et al., 2016; Chen et al., 2010; C. E. Kim et al., 2010; Nery et al., 2011; Rittiner et al., 2016; Zhao et al., 2016). Because many of these defects are apparent only following stress treatment, we additionally performed the proteomic experiment in samples after six hours of thapsigargin (Tg) exposure, a compound that causes cell stress by releasing internal calcium stores. With Tg treatment, we found that cytosolic samples showed a similar number of genotype-dependent differences as non-stressed (Veh) samples (115 proteins) (Fig. 2C). However, in striking contrast to cytosolic fractions, Tg treatment caused a greater than 3-fold increase in genotype-dependent differences in the nuclear proteome (624 proteins) relative to the non-stressed samples (187 proteins) (Fig. 2D). These data reveal that among cytosolic and nuclear compartments in basal and stressed states, the DYT-TOR1A genotype most severely disrupts the composition of the nuclear proteome and does so selectively in the presence of a cellular stressor.

Given the similarity in the number of protein differences between the cell stress and basal conditions in the cytosolic fractions, we next examined the extent to which genotype-dependent differences were due to the same proteins being affected in multiple conditions. Surprisingly, there was little overlap - with less than 10% of proteins being shared between the stressed and basal state conditions (Fig. S2B). These findings indicate that, in both the nuclear and cytosolic fractions, the proteome disruptions caused by DYT-TOR1A under cell stress affects proteins that are largely distinct from those in the basal state.

Next, we evaluated genotype-dependent differences in the subcellular fractionation of TorsinA itself. Multiple prior studies have found that the DYT-TOR1A mutation of *Tor1a* (ΔE TorsinA) drives TorsinA mislocalization from the ER to the nuclear envelope (Bragg et al., 2004; Calakos et al., 2010; Gonzalez-Alegre & Paulson, 2004; Goodchild & Dauer, 2004; Hewett et al., 2000; Kustedjo et al., 2000; Liang et al., 2014; Naismith et al., 2004; Torres et al., 2004). The results for TorsinA were not included in our proteomic analysis because TorsinA was identified by only a single unique identifying peptide, and this is associated with an increased risk for misidentifying proteins (Carr et al., 2004). However, given the particular relevance of TorsinA data to this study, we used the single peptide data to examine its distribution between nuclear and cytosolic compartments. We found that relative to WT samples, TorsinA levels were significantly lower in cytosolic fractions of DYT-TOR1A samples (p = 0.004) and that there was also a non-significant trend toward higher levels of TorsinA in nuclear fractions of DYT-TOR1A samples (p = 0.232) significantly altering its distribution to be biased towards the nuclear compartment (N/C ratio) (Fig. 2E).

Interpretation of data from our approach and most other subcellular proteomic approaches relies upon the assumption that cell constituents fractionate normally (Ortega et al., 2020; Tribl et al., 2005). In Figure 1B-C, we observed that there were no obvious solubilization differences between WT and DYT-TOR1A MEFs. Nonetheless, for any specific protein of interest, the use of an orthogonal methodology would be a desirable validation step. To date, conventional immunofluorescence has not revealed the TorsinA redistribution in genetic construct-valid DYT-TOR1A cells that we detected here using quantitative proteomics. To better understand the misdistribution of ΔE TorsinA and address the integrity of nuclear envelope partitioning in DYT-TOR1A cells, we evaluated the partitioning of known nuclear envelope proteins, the LINC complexes in WT and DYT-TOR1A MEFs (Fig. S4). We found that LINC proteins present in our datasets enriched in the nuclear fraction as expected and partitioned similarly in WT and DYT-TOR1A samples. We also present two examples of conventional ICC validation. Levels of Tbce and Pds5b were significantly altered in DYT-TOR1A in the nuclear compartment under Tg cell stress condition (Fig. S5-S6). Proteomic analysis of Tbce showed a compartment-specific decrease in the Tg-nucleus and no difference in cytosolic levels (Fig. S5B, D); findings which were replicated with conventional ICC (Fig. S5C, E). Pds5b levels were significantly increased in DYT-TOR1A Tg-nuclear samples (Fig. S6B).

Using ICC, nuclear Pds5b immunostaining intensity was significantly different by genotype; however, instead of increased, Pds5b staining was significantly decreased in DYT-TOR1A (Fig. S6C). Interestingly, in WT cells, Pds5b was more commonly in strongly staining puncta, raising the possibility that reduced solubility of punctate Pds5b might give rise to the proteomic result of lower levels in WT cells (Fig. S6D, E). To summarize, while ICC for both proteins confirmed genotype effects, these two examples highlight the range of disruptions that might underlie the proteomic results.

### Thapsigargin Stress-Responsive Proteins in WT MEFs

Thus far, our proteomic analyses reveal that the largest DYT-TOR1A genotype-dependent disruption to the proteome was observed in the nuclear compartment under cell stress. Since a number of prior studies have shown abnormalities in cell stress responses in DYT-TOR1A models, we asked whether the nuclear proteomic disruptions were predominantly composed of proteins whose levels were normally modulated by cell stress. To address this, we first used the WT datasets to identify the normal subset of stress-responsive proteins – i.e. proteins whose levels significantly changed in response to thapsigargin cell stress. For each protein and subcellular fraction of the WT samples, the ratio of levels in the Tg and Veh conditions were calculated. Using thresholds of ±1.5 FC and p-value < 0.05, we identified a total of 513 proteins that we hereafter refer to as the “stress-responsive proteins”.

Consistent with the global reduction in protein synthesis rates that occurs following ER stress (Ron, 2002), Tg cell stress tended to downregulate more proteins (373 proteins) than it upregulated (140 proteins) (Fig. 3A-B). Among the upregulated stress-responsive proteins in WT samples, GO analysis revealed significant enrichment for proteins associated with the PERK-mediated unfolded protein response (p=0.002, Fold Enrichment = 10.8). This enrichment is expected given that thapsigargin is thought to promote cellular stress response through a PERK-dependent mechanism (H. P. Harding et al., 2000). Individual examples of two proteins associated with this pathway, Herpud1 and NFκB p105, are shown (Fig. 3C-D).

**Fig. 3.**
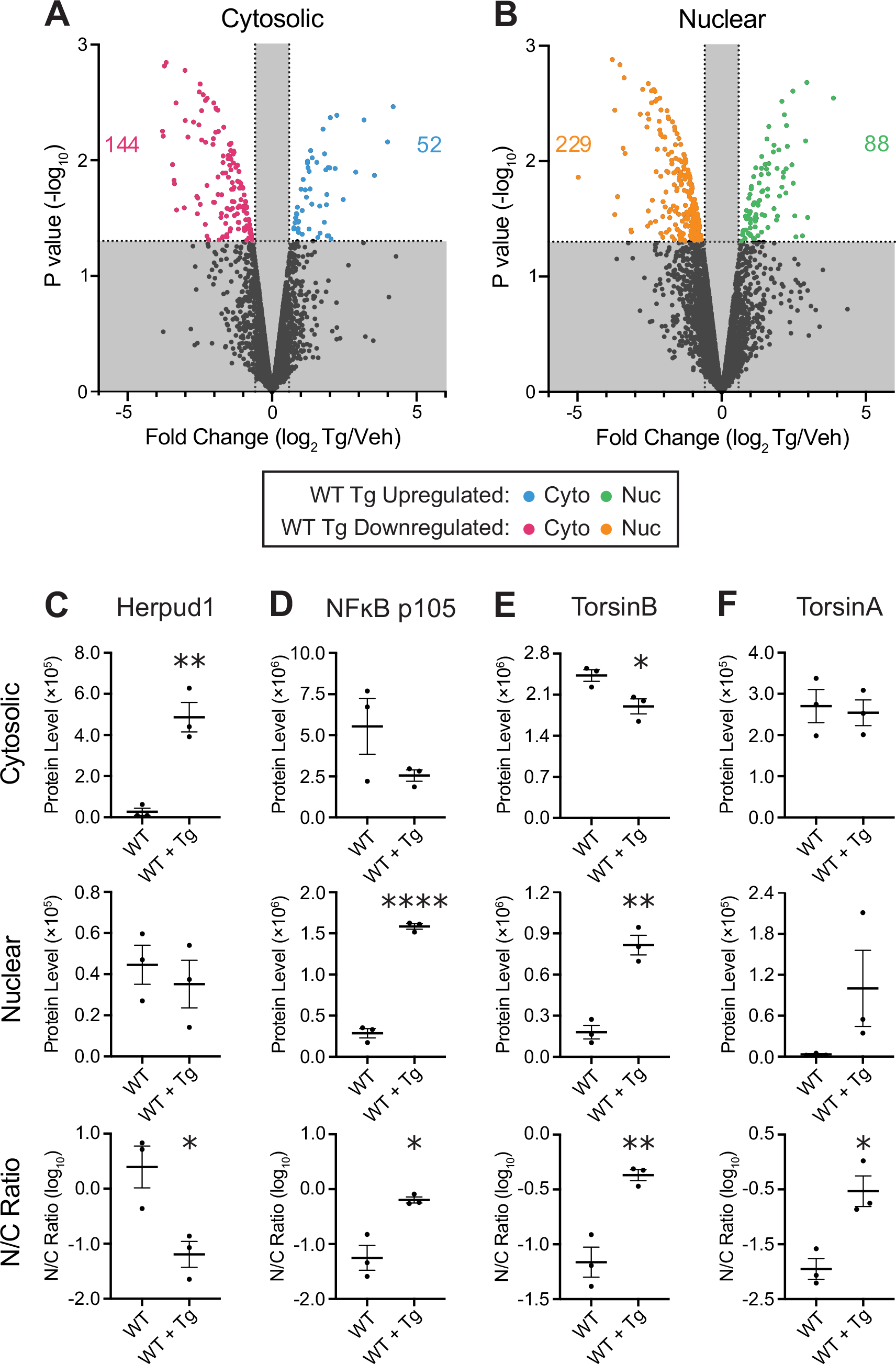
Thapsigargin stress modulation of protein levels in wildtype MEFs. (A-B) Volcano plots showing proteome-wide differences in protein abundances due to thapsigargin (Tg) treatment in wildtype cytosolic (A) and nuclear (B) fractions. Differences in protein abundance are represented as fold change (using log2 transform) and p-value is calculated by unpaired t-test for each protein (n=3 biological replicates). Horizontal dashed line indicates p-value less than 0.05. Vertical dashed lines indicate fold changes of ±1.5 fold change (FC) (C-F) Cytosolic and nuclear fraction protein levels (a.u.) and the corresponding nuclear:cytosolic (N/C) ratios are shown for Herpud1, NFκB p105, TorsinB, and TorsinA in Veh (WT) and Tg treatment (WT + Tg) conditions (n = 3 biological replicates; *p<0.05;**p<0.01;****p<0.0001, unpaired t-test). Error bars indicate S.E.M..

Lastly, in reviewing the stress-responsive proteins we identified in WT samples, we were surprised to see the paralog of TorsinA, TorsinB, among them. TorsinB levels were significantly increased by stress in the nuclear fractions and decreased in the cytosolic fractions, resulting in a greater than doubling of its relative nucleocytoplasmic distribution (nuclear:cytosolic protein levels = N/C ratio) (Fig. 3E). A similar trend was seen for TorsinA (Fig. 3F). This finding suggests for the first time that redistribution of Torsin proteins toward the nucleus may be part of the normal cellular stress response.

### DYT-TOR1A MEFs Show Basal Elevations and Blunted Stress Responses of Normally Stress-Responsive Proteins

Having defined the normal stress-responsive shifts in the proteome, we next evaluated whether the WT stress-responsive proteins were enriched among DYT-TOR1A disrupted proteins. We found that stress-responsive proteins were significantly enriched among the DYT-TOR1A disrupted proteins. This enrichment was present across all conditions and subcellular fractions. To visualize this, in Fig. 4 we show the DYT-TOR1A/WT comparison datasets using the color scheme from Fig. 3 to indicate the nature of the WT stress response (i.e. upregulated or downregulated in Fig. 3A-B). Notably, under basal conditions (Veh), WT stress-responsive proteins were not uniformly distributed but rather tended to align with the directionality of their normal modulation by cell stress. As a group, the proteins upregulated by thapsigargin within WT MEFs were significantly enriched among the subset of proteins upregulated in DYT-TOR1A basally (Cyto: p=7.68e-8; Nuc: p=5.98e-12) (Fig. 4A-B). Similarly, the group of proteins downregulated by Tg treatment in WT MEFs were significantly enriched among the subset of proteins downregulated in DYT-TOR1A basally (Cyto: p=6.92e-6; Nuc: p=8.81e-11) (Fig. 4A-B). This analysis reveals that the basal state proteome of DYT-TOR1A MEFs reflects an activated cell stress state prior to exogenous Tg cell stress treatment.

**Fig. 4.**
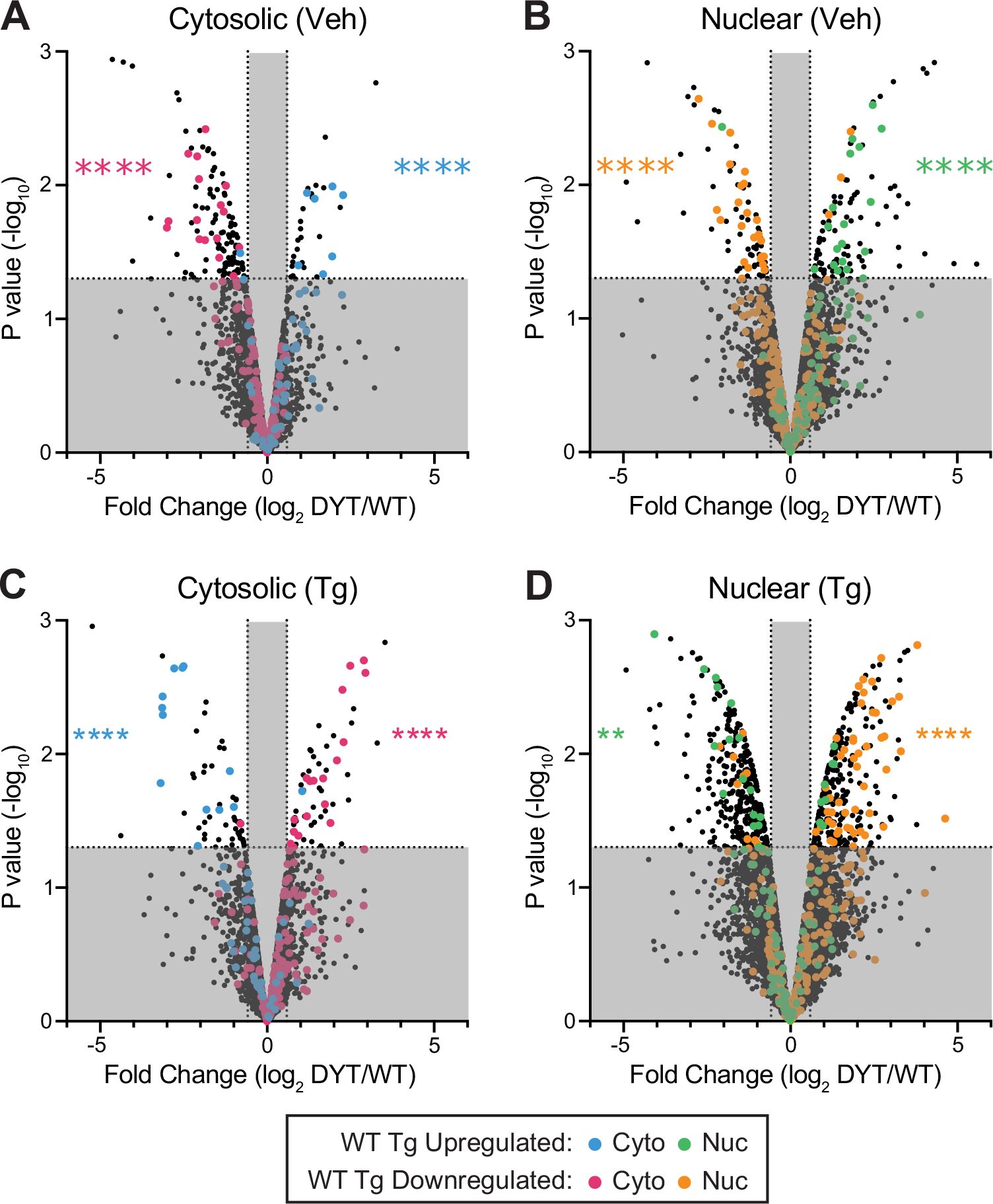
WT Tg-stress-responsive proteins are non-randomly distributed among DYT-TOR1A genotype-dependent proteome disruptions. Volcano plots showing proteome-wide differences in protein abundances between DYT-TOR1A and WT cytosolic and nuclear fractions under basal (A, B) and thapsigargin-treated (C, D) conditions. For proteins found to be Tg-modulated in WT samples (Fig. 3), the directionality of the WT sample modulation is indicated by symbol color (see Legend). Differences in protein abundance are represented as fold change (using log2 transform) and p-value is calculated by unpaired t-test for each protein (n=3 biological replicates). Horizontal dashed line indicates p-value less than 0.05. Vertical dashed lines indicate fold changes of ±1.5 FC. Asterisks indicate p-value calculated by Fisher’s exact test for overlap between the indicated subsets of DYT1 disrupted proteins and WT Tg-stress-responsive proteins in the white quadrants (**p<0.01; ****p<0.0001).

Following Tg stress treatment, we again found significant overlap between the normal (WT) stress-responsive proteins and DYT-TOR1A genotype-dependent protein disruptions in both subcellular compartments (Fig. 4C-D). However, under Tg cell stress, the directionality of the genotype-dependent disruptions was opposite to that of the normal stress response (visualized by the colored symbols concentrating on the opposite side of the volcano plots, Fig. 4C-D). Upregulated WT stress-responsive proteins were significantly enriched among the subset of downregulated proteins in Tg-treated DYT-TOR1A fractions when compared to Tg-treated WT fractions (Cyto: p=4.38e-12; Nuc: p=9.92e-3) (Fig. 4C-D). Likewise, downregulated WT stress-responsive proteins were significantly enriched among the subset of upregulated proteins in Tg-treated DYT-TOR1A fractions compared to Tg-treated WT fractions (Cyto: p=4.67e-13; Nuc: p=2.09e-12) (Fig. 4C-D). These findings are consistent with a blunted stress response in DYT-TOR1A MEFs following Tg cell stress.

At the level of individual proteins, we noticed that these same trends could be seen in the N/C ratios of NFκΒ p105 and TorsinA, but also, that for others, stress-regulation was unaffected by the DYT-TOR1A genotype (e.g. Herpud1 and TorsinB) (Fig. 5A-D, S3). Given this variation, we sought to quantify the average degree of modulation across the entire population of WT stress-responsive proteins. We measured the fold change for each of the 140 WT stress-upregulated proteins and 373 WT stress-downregulated proteins in each condition relative to its level in the basal state WT samples (Veh), and then calculated the mean fold change for all up- or downregulated proteins in each condition (Fig. 5E-F). This analysis shows that in the basal state, levels of WT stress-responsive proteins in DYT-TOR1A samples already have modulations in the direction consistent with stress responses – the mean level of WT stress-upregulated proteins was ∼60 percent (p = 8.63e-19) higher than WT basal levels, while the mean level of WT stress-downregulated proteins was ∼15 percent lower (p = 1.95e-5). These findings indicate that the nuclear and cytosolic proteomes in DYT-TOR1A MEFs show modulations consistent with a stressed state basally.

**Fig. 5.**
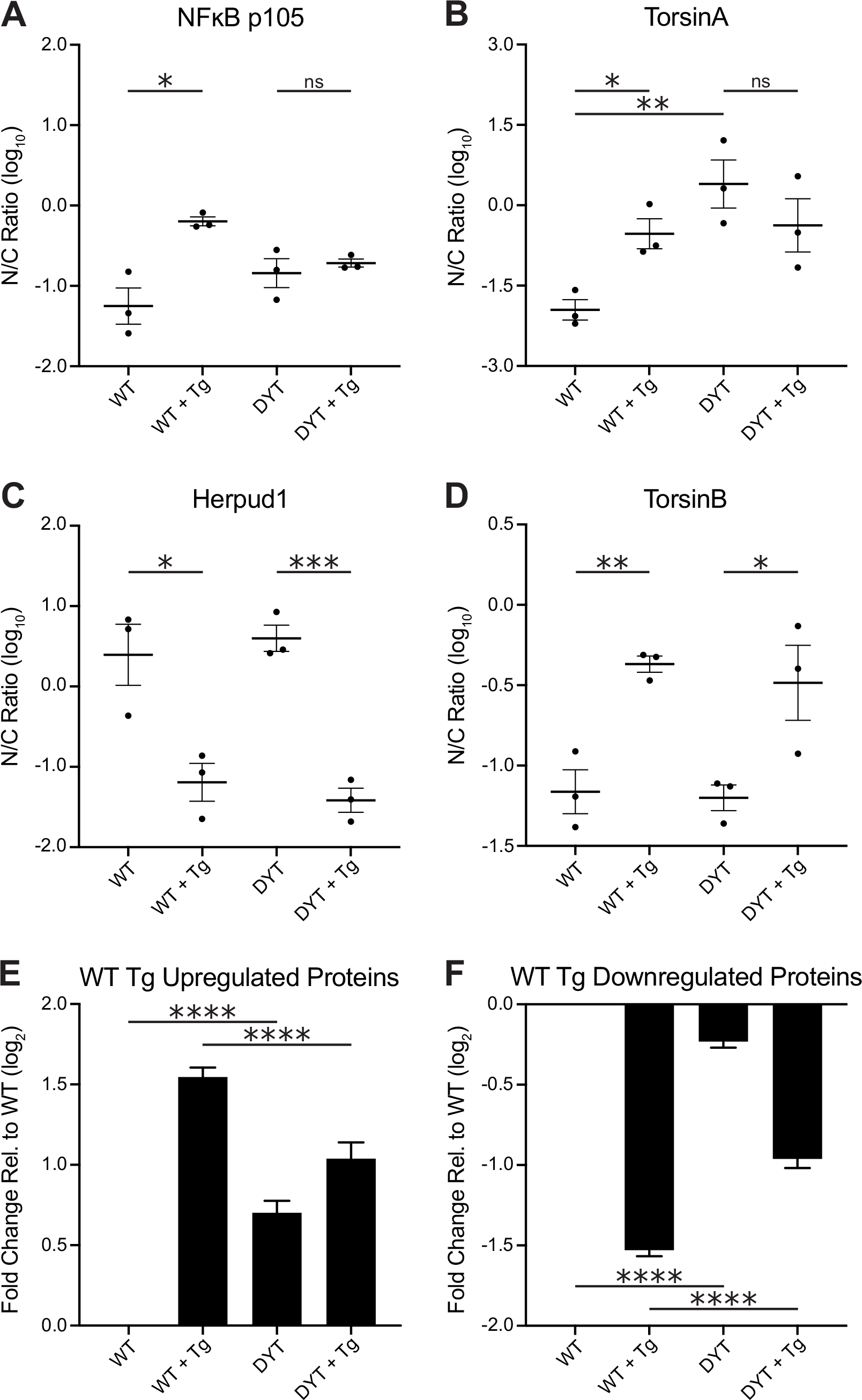
DYT-TOR1A MEFs show proteome-wide disruptions consistent with an elevated basal-state stress response and a blunted response to Tg stress. (A-D) Individual protein examples of Tg stress-induced shifts in protein abundance between nuclear and cytosolic fractions are shown for NFκB p105 (A), TorsinA (B), Herpud1 (C), and TorsinB (D) in WT and DYT-TOR1A MEFs (n=3 biological replicates) (*p<0.05; **p<0.01; ***p<0.001, unpaired t-test). (E-F) Mean fold change calculated across all WT Tg-stress-responsive upregulated (n=140) (E) and downregulated (n=373) (F) proteins. For this calculation, protein levels (a.u.) from WT and DYT-TOR1A fractions treated with either Tg or Veh were normalized to the protein level measured in the WT Veh-treated fraction from the same subcellular compartment (n=3 FC values per protein to calculate mean value). Significance was determined by unpaired t-test between mean FC values for 140 upregulated stress-responsive proteins (E)(****p<0.0001) and 373 downregulated stress-responsive proteins (F)(****p<0.0001). Error bars indicate S.E.M..

In WT cells, cell stress by Tg exposure caused a threefold change in the mean level of upregulated and downregulated WT stress-responsive proteins (Fig. 5E-F). By comparison, in DYT-TOR1A MEFs, Tg caused only a twofold change in the levels of these same WT stress-responsive proteins (upregulated: p= 8.63e-19; downregulated: p=3.40e-16) (Fig. 5E-F). These results indicate that while the DYT-TOR1A proteome does respond to cellular stress, the magnitude of the response is significantly reduced.

### Dysregulation of WT Stress-Responsive Proteins Does Not Explain Accentuated Proteome Disruption in Stressed Nuclear DYT-TOR1A Fractions

Thus far, we have found that WT stress-responsive proteins are significantly enriched among the proteins whose levels are disrupted by DYT-TOR1A (Fig. 4). Because this enrichment is similarly observed in both subcellular compartments and under both basal and Tg cell stress (Fig. 4), it cannot explain the large, Tg and nucleus-selective disruption of 624 proteins caused by DYT-TOR1A. Moreover, stress-responsive proteins explain only 13% of the total number of disrupted proteins in the Tg nuclear samples (Fig. 6A). We therefore sought to further understand the nature of the proteins disrupted by DYT-TOR1A in the nucleus under stress (Fig. 6A-C). A GO analysis was performed on the set of 624 proteins differentially regulated in DYT-TOR1A + Tg nuclear fractions as compared to the WT + Tg nuclear fractions (Fig. 6D, Table S1). Enrichments included “neuron cellular homeostasis” (p=0.002, Fold Enrichment=4.5), “mRNA-containing ribonucleoprotein (RNP) complex export from the nucleus” (p=0.004, Fold Enrichment = 1.8), and “axonal transport” (p=0.006, Fold Enrichment=2.3) (Fig. 6D and Table S1). These GO terms suggest that in addition to the pervasive disruptions in cell stress responses, the DYT-TOR1A genotype may also cause particular liabilities in the nucleus under cell stress among proteins generally important for synaptic and neuronal function.

**Fig. 6.**
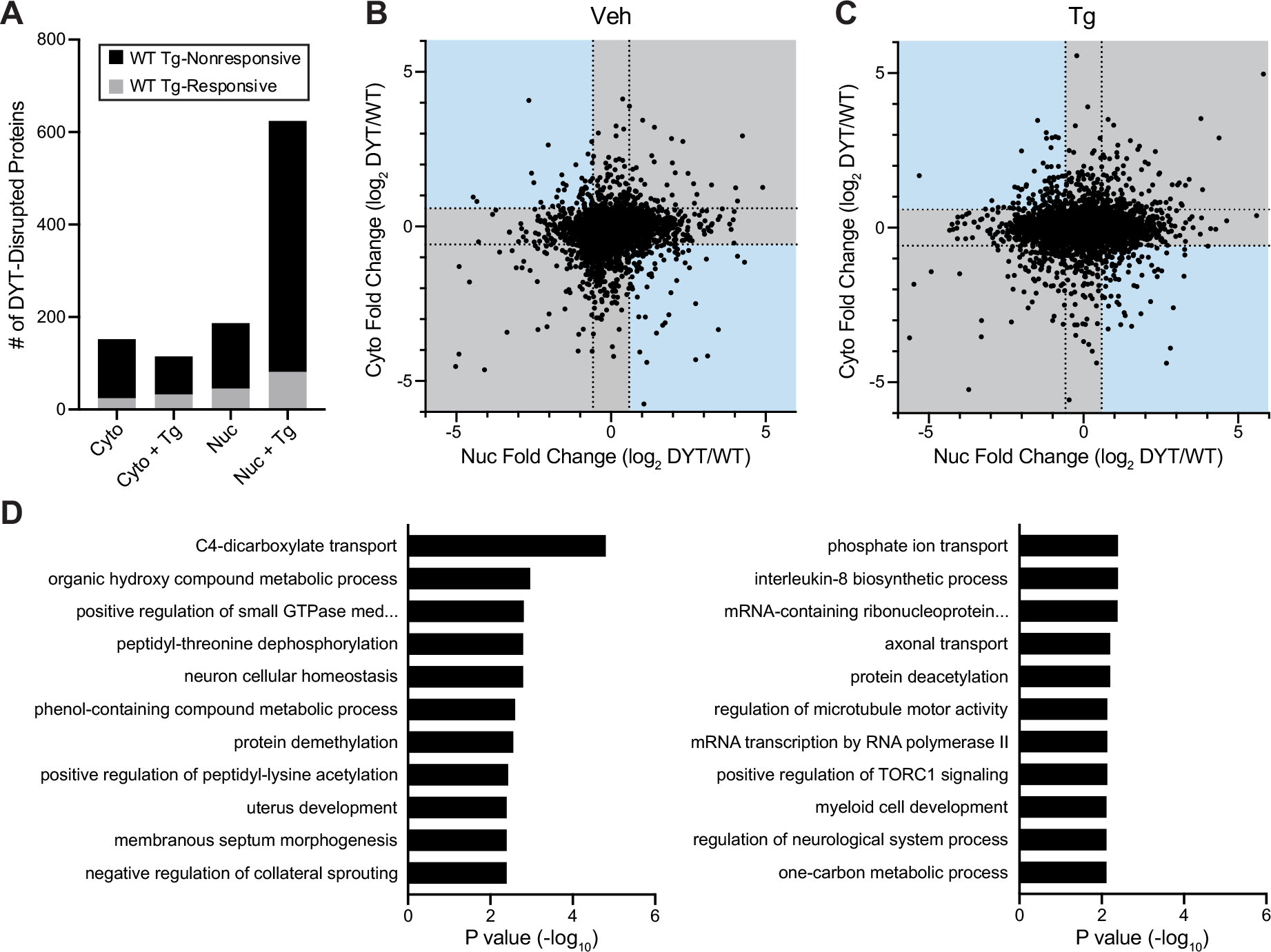
Stress-dependent nuclear proteome disruptions in DYT-TOR1A MEFs associate with critical neuronal processes. (A) Absolute number of proteins with significant genotype effects (FC > ±1.5, p<0.05) in each cellular fraction and treatment condition. (B, C) Proteome-wide results for genotype effects comparing directionality of fold change between the nuclear and cytosolic compartments in the vehicle (B) and thapsigargin (C) conditions. Blue shading denotes zones of reciprocal relationships. Gray shaded regions are FC less than ±1.5. (D) Gene Ontology analysis of the 624 proteins in the Tg-nuclear fractions with significant DYT-TOR1A genotype-dependent effects (FC > ±1.5, p<0.05).

## Discussion

In this study, we performed quantitative proteomic analysis of nuclear and cytosolic-enriched fractions prepared from DYT-TOR1A and WT MEFs in order to identify cell compartment-specific disruptions. This proteomic approach was driven by our hypothesis that some protein level disruptions caused by the DYT-TOR1A genotype may differentially manifest between the nuclear and cytosolic compartments given associations of TorsinA and ΔE TorsinA with nuclear envelope structure and nuclear transport functions (Chase et al., 2017; Ding et al., 2021; Gonzalez-Alegre & Paulson, 2004; Goodchild et al., 2005; Goodchild & Dauer, 2004, 2005; György et al., 2018; Jokhi et al., 2013; M. T. Jungwirth et al., 2011; Laudermilch et al., 2016; Naismith et al., 2004; Nery et al., 2011; Pappas et al., 2018; Rampello et al., 2019; Tanabe et al., 2016; VanGompel et al., 2015). To date, proteomic studies of dystonia have examined whole cell and tissue lysates (Beauvais et al., 2016; Martin et al., 2009; Zakirova et al., 2018). Using nucleocytoplasmic proteomics, we discovered that the DYT-TOR1A mutation disproportionately causes the nuclear proteome to be disrupted by cell stress. There were 3-fold more disrupted proteins in the thapsigargin-treated nuclear proteome than any other condition (Fig. 6). Although this effect occurred under Tg cell stress and we found that levels of many proteins that were normally modulated by Tg were disrupted in DYT-TOR1A, the majority of the affected proteins were not part of the WT stress-responsive subset. In addition to this novel finding, our datasets provide further support for several prior observations in the field. For example, in both subcellular compartments, DYT-TOR1A samples show evidence of impaired mitochondrial function and stress responses (Beauvais et al., 2016; Cao et al., 2005; Chen et al., 2010; Martin et al., 2009; Nery et al., 2011; Rittiner et al., 2016).

To begin to address the potential impact of the stress-dependent nuclear proteome disruptions in DYT-TOR1A MEFs on dystonia pathophysiology, we conducted Gene Ontology analysis. Despite the source cells being non-neuronal, a number of GO terms with particular importance for neuronal function were enriched (Fig. 6). These include: “Neuron Cellular Homeostasis”, “Negative Regulation of Collateral Sprouting”, “mRNA-containing Ribonucleoprotein Complex Export from the Nucleus”, “Axonal Transport”, “Regulation of Microtubule Motor Activity”, “Positive Regulation of TORC1 Signaling”, and “Regulation of Neurological System Process”. One of the identified biological functions enriched among the stress-dependent nuclear proteome disruptions caused by the DYT-TOR1A genotype was “mRNA-containing ribonucleoprotein complex export from the nucleus”. Prior studies have made an association between a particular type of RNP complex, a “megaRNP” and TorsinA function (Jokhi et al., 2013). In *D. melanogaster*, nuclear egress of the megaRNP is essential for proper synaptic development of the neuromuscular junction (Speese et al., 2012). More generally, ribonucleoprotein complexes are known to play an integral role in the maintenance, stress response, and plasticity of synapses, and RNP disruptions are associated with a number of neurological diseases (Khalil et al., 2018; Ross Buchan, 2014). Therefore, taken together, the GO processes involving RNP complexes, transport processes, and neuronal homeostasis are potentially interrelated and predict impact on critical brain processes. Our findings suggest that these processes may be most disrupted in settings of cell stress - whether due to inherent states, such as development, or exogenous cell stressors. An important future direction is to determine whether the stress-dependent vulnerability of the nuclear proteome observed in this study also exists within the mammalian central nervous system and, if so, which neural cells are most affected by this vulnerability.

Our proteomic data identify protein levels that differ because of the DYT-TOR1A genotype. In interpreting these data, there are a number of factors that could lead to altered protein levels. Such factors include changes in protein synthesis rates, protein degradation rates, transport between subcellular compartments or aggregation state. TorsinA is well known to move between the lumen of the endoplasmic reticulum and the outer nuclear envelope in an ATP-hydrolysis dependent manner (Goodchild & Dauer, 2004, 2005; Naismith et al., 2004; Vander Heyden et al., 2009; Zhao et al., 2013). Moreover, a number of experimental approaches have shown that TorsinA deletion or ΔE TorsinA overexpression disrupts this trafficking and nuclear envelope structure (Ding et al., 2021; Gonzalez-Alegre & Paulson, 2004; Goodchild et al., 2015; Goodchild & Dauer, 2004; M. T. Jungwirth et al., 2011; Naismith et al., 2004; Torres et al., 2004; Vander Heyden et al., 2009). We therefore found it notable that our proteomic data did not provide support for a model where ΔE TorsinA restricts protein trafficking between the nucleus and cytosol. More specifically, we performed an analysis across the entire detected proteome irrespective of the p-values to determine if we could detect even a trend for proteins to show reciprocal relationships between nuclear and cytosolic fractions (i.e. lowering in one compartment and increasing in the other) (Fig. 6B-C, blue areas). Such changes would be represented in the colored quadrants of those graphs and create an elliptical skewing. However, in this analysis, we saw no trends for reciprocal changes. Instead, we see that basally, there was little if any skew (Fig. 6B) and that with thapsigargin, the major modulation was restricted to the nuclear compartment (e.g. data expanding horizontally, nuclear fold-change axis) with little change vertically (cytosolic fold-change axis) (Fig. 6C). We therefore favor models other than nucleocytoplasmic transport defects to explain the bulk of proteome disruption in DYT-TOR1A MEFs. As one example of alternative models, compartment-specific protein level disruptions could arise by dysregulation of compartment-specific protein degradation mechanisms, such as the nuclear proteasome or ER-associated degradation mechanisms (Enenkel, 2014; Nery et al., 2011). We are interested in testing this possibility in future studies.

In addition to identifying nuclear compartment-specific disruptions that were largely unrelated to WT stress-responsive proteins, we also made a number of novel observations about the integrity of the stress response in DYT-TOR1A cells. Foremost among these, by using a proteome-wide approach as opposed to monitoring a small number of proteins of interest, we found that, as a group, proteins whose levels were normally modulated by cell stress (in this experiment, by thapsigargin) tended to show deviations toward their stress response in the basal state in DYT-TOR1A cells (Fig. 4,5E-F). This result suggests that the DYT-TOR1A genotype induces a basally elevated stress state.

Basal elevation of the cellular stress response has been observed in other DYT-TOR1A cellular models. For example, BiP, a key stress-responsive protein, is upregulated in an unstressed *C. elegans* DYT-TOR1A transgenic model (Chen et al., 2010). In this study, the investigators further noted that in response to cell stress (using tunicamycin, a glycosylation inhibitor), DYT-TOR1A model worms had an exaggerated BiP response. Tunicamycin increased BiP expression in both WT and DYT-TOR1A samples, but to a greater degree in DYT-TOR1A. To explain the elevated stress response both basally and following stress, the authors speculated that DYT-TOR1A cells have a reduced buffering capacity against cell stress. In our data, we confirm the observations of Chen and colleagues – levels of BiP are greater in DYT-TOR1A than WT basally and increase more in DYT-TOR1A than WT in response to thapsigargin (Fig. S7B). However, by looking at the entire set of experimentally identified stress-responsive proteins, we further recognized that the BiP response was not representative of the average response to cell stress in DYT-TOR1A MEFs. Instead, we find that the DYT-TOR1A genotype *lowers* the overall stress response relative to WT (Fig. 5E-F). Blunting of the stress response has also been reported in primary fibroblasts from DYT-TOR1A patients and cerebellar tissue from DYT-TOR1A mouse models (Beauvais et al., 2016; Rittiner et al., 2016). Based on our new findings, we postulate that homeostatic dysregulation of stress signaling pathways may have developed due to the chronically elevated stress response in DYT-TOR1A cells that exists prior to exogenous stress treatment. An important area for future studies in DYT-TOR1A is to identify the biological mechanisms inciting basal cell stress and driving homeostatic dysregulation.

Although our proteomic experiments were not designed to address single protein-level hypotheses, our results make three preliminary novel observations regarding Torsins. First, to our knowledge, we make the first observation that TorsinB is a stress-responsive protein. Second, levels of TorsinA also appear to be stress-modulated. However, in contrast to TorsinB, TorsinA stress modulation is impaired (occluded) in DYT-TOR1A cells, while the response of TorsinB appears normal (Fig. 5B, D). This difference between TorsinA and TorsinB is noteworthy because a number of prior studies have highlighted the potential for TorsinB to substitute for loss of TorsinA function in DYT-TOR1A (M. Jungwirth et al., 2010; Tanabe et al., 2016; Vasudevan et al., 2006), including a recent study which shows that overexpression of TorsinB rescues abnormal movement phenotypes observed in forebrain specific *Tor1a* and *Tor1a/Tor1b* combined conditional knockout as well as selective *Tor1a^Δ^*^GAG^ knock-in mouse models (Li et al., 2020). Our data provide additional support for the rationale of such therapeutic approaches. Third and last, with the sensitivity afforded by proteomic methodologies, we find that TorsinA is mislocalized toward the nuclear compartment in DYT-TOR1A cells with genetic construct validity. Although mislocalization of ΔE TorsinA has been widely observed across labs and experimental settings (Bragg et al., 2004; Calakos et al., 2010; Gonzalez-Alegre & Paulson, 2004; Goodchild & Dauer, 2004; Hewett et al., 2000; Kustedjo et al., 2000; Liang et al., 2014; Naismith et al., 2004; Torres et al., 2004), to our knowledge it has never been documented in a construct-valid genetic model for DYT-TOR1A dystonia (e.g. *Tor1a* ^ΔGAG*/+*^). Our results therefore provide experimental support for the idea that ΔE TorsinA mislocalization exists in the genetically relevant setting and it may only be the matter of degree that differs from overexpression models.

## Conclusions

We have newly identified a stress-dependent and nuclear compartment-specific proteome disruption caused by the DYT-TOR1A genotype. Our findings suggest that key brain processes involving neuronal homeostasis, transport, and RNP export may be selectively impaired in DYT-TOR1A by cell stress through effects on the nuclear proteome. Alongside further understanding TorsinA’s role in modifying cellular stress responses, our results open a new research direction for DYT-TOR1A dystonia pathophysiology which is to understand this compartment-specific vulnerability and its consequences for brain function.

Author Contributions:

KS – Conceptualization, Investigation, Formal analysis, Writing – Original Draft, Review & Editing

ZFC –Supervision, Validation, Writing – Review & Editing

NC – Conceptualization, Funding acquisition, Supervision, Writing – Review & Editing

## Acknowledgements

The authors wish to acknowledge critical expertise and suggestions provided by Shataakshi Dube, William Dauer, Connor King, Miranda Shipman, Erik Soderblom and the Duke Proteomics and Metabolomics Core. K.S. thanks the members of his undergraduate thesis committee, Ron Grunwald and Matthew Oliver, for continued support and guidance.

## Abbreviations

Cyto: cytosolic fraction
DYT: DYT-TOR1A genotype
FC: fold change
GO: Gene Ontology
LC: liquid chromatography
MEF: mouse embryonic fibroblast
MS: mass spectroscopy
Nuc: nuclear fraction
RNC: relative nuclear concentration
RNP: ribonucleoprotein
Tg: thapsigargin
Veh: vehicle control
WT: wildtype

---

**Fig. S1.**
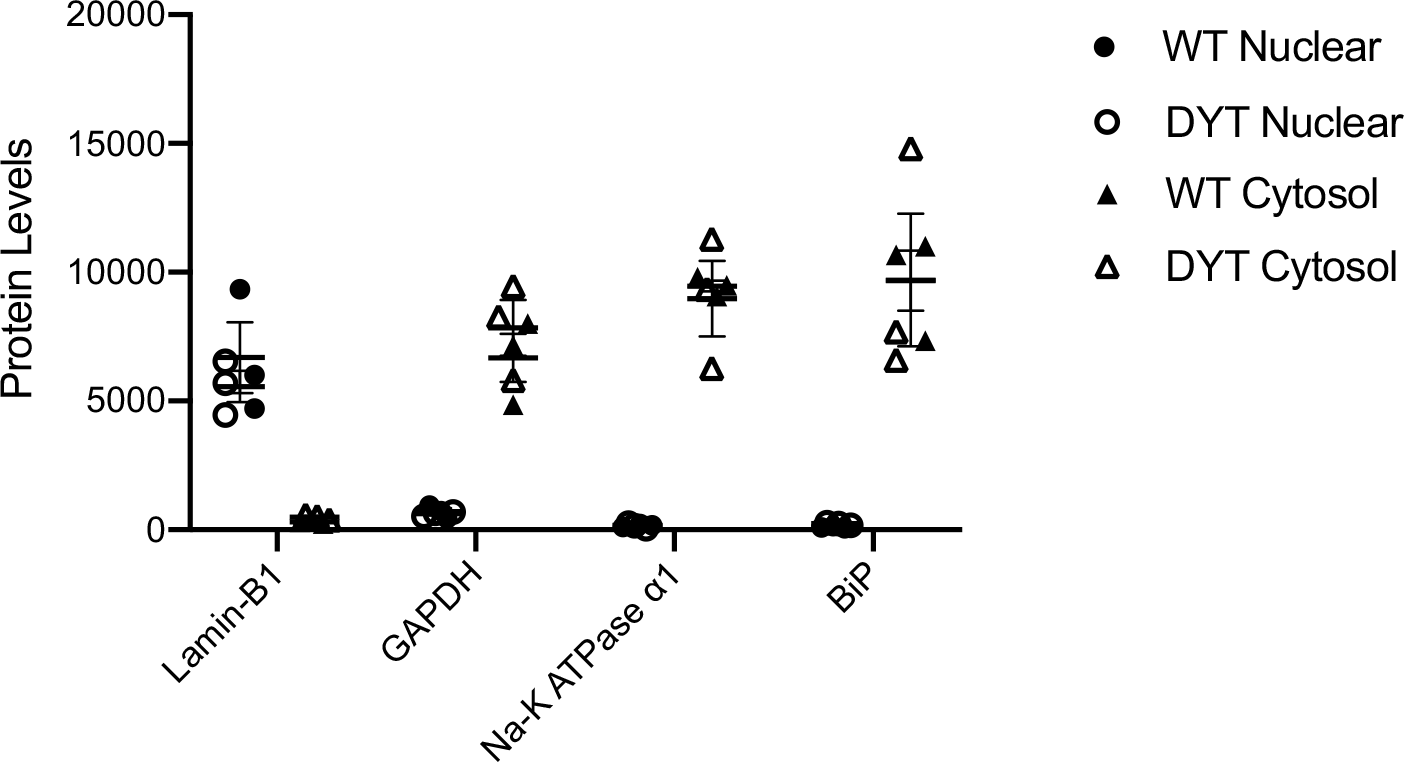
Quantification of Western blot of WT and DYT-TOR1A samples under basal conditions. Western blot protein level (a.u.) was quantified by densitometric analysis of the fluorescent signal. N = 3 independent measures per condition.

**Fig. S2.**
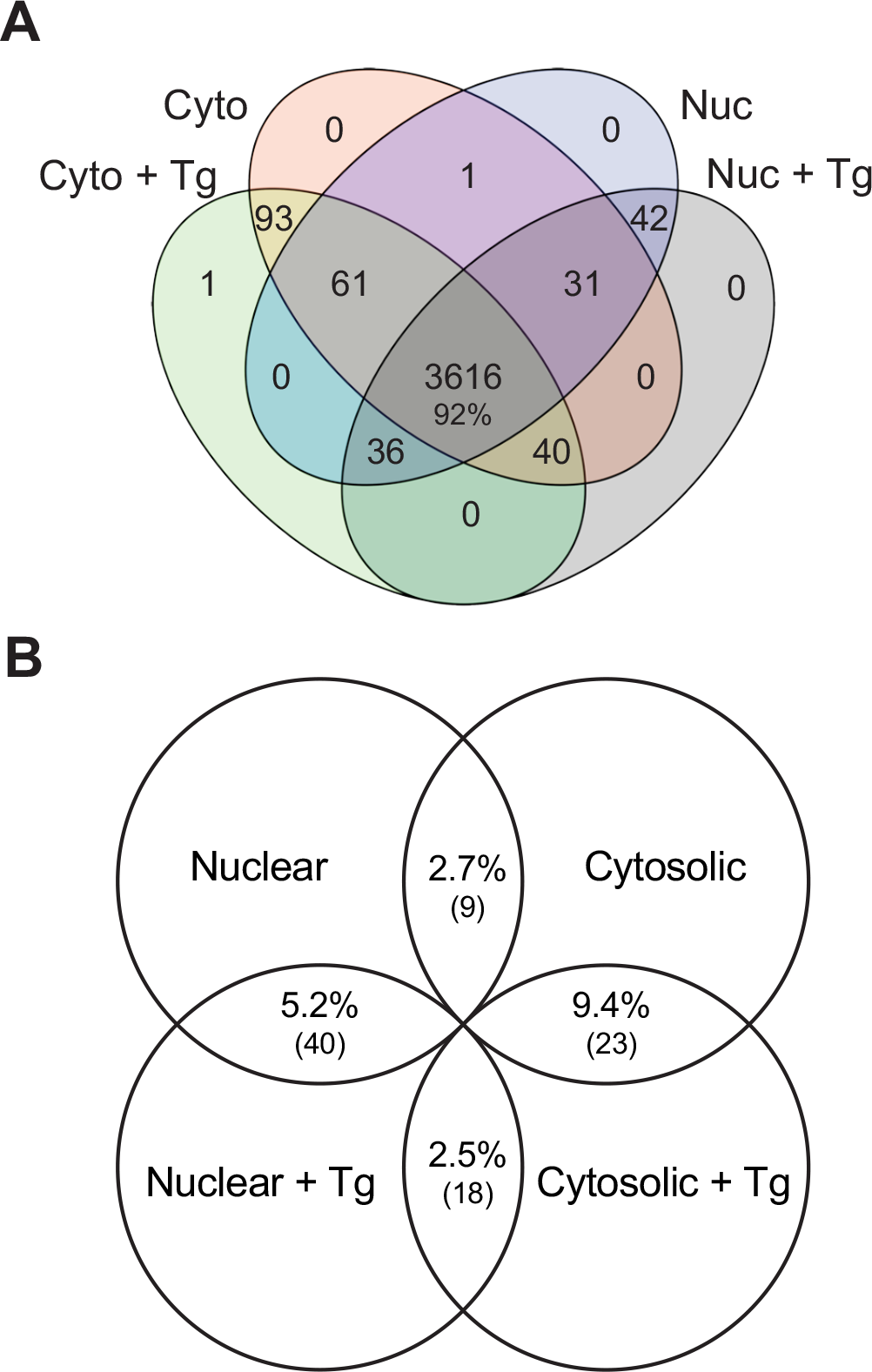
Overlap of LC/MS/MS identified proteins across experimental conditions. (A) Overlap of LC/MS/MS identified proteins between nuclear and cytosolic fractions treated with either Tg or Veh. (B) Pairwise overlap of DYT-disrupted proteins between nuclear and cytosolic fractions treated with either Tg or Veh.

**Fig. S3.**
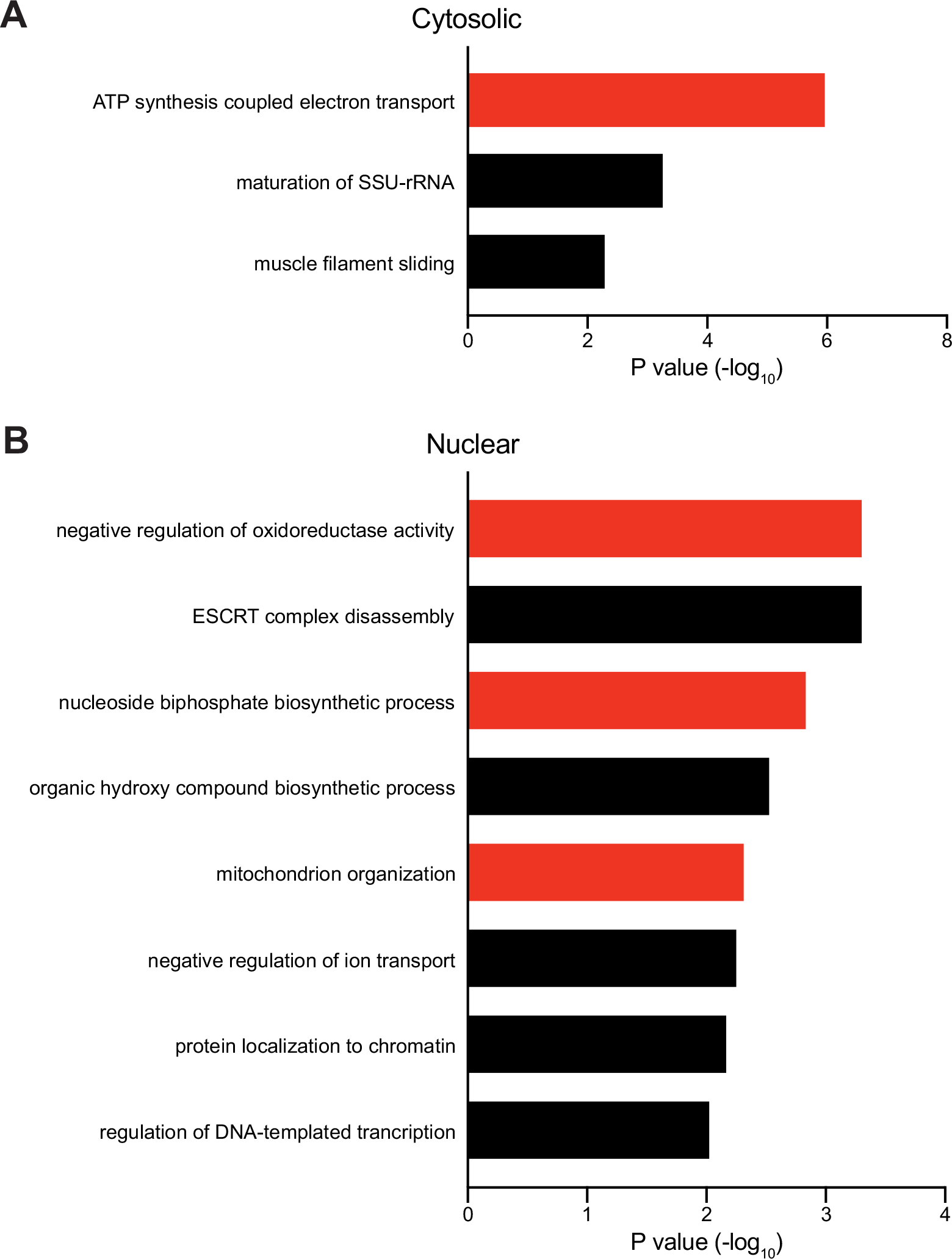
Gene Ontology analysis of DYT-TOR1A genotype-dependent proteome disruptions. (A) Gene Ontology analysis of the 152 cytosolic fraction proteins showing DYT-TOR1A/WT or WT/DYT-TOR1A FC > 1.5 and p < 0.05. (B) Gene Ontology analysis of the 187 nuclear fraction showing DYT-TOR1A/WT or WT/DYT-TOR1A FC > 1.5 and p < 0.05. Red bars highlight Gene Ontology terms associated with mitochondrial organization or ATP metabolism.

**Fig. S4.**
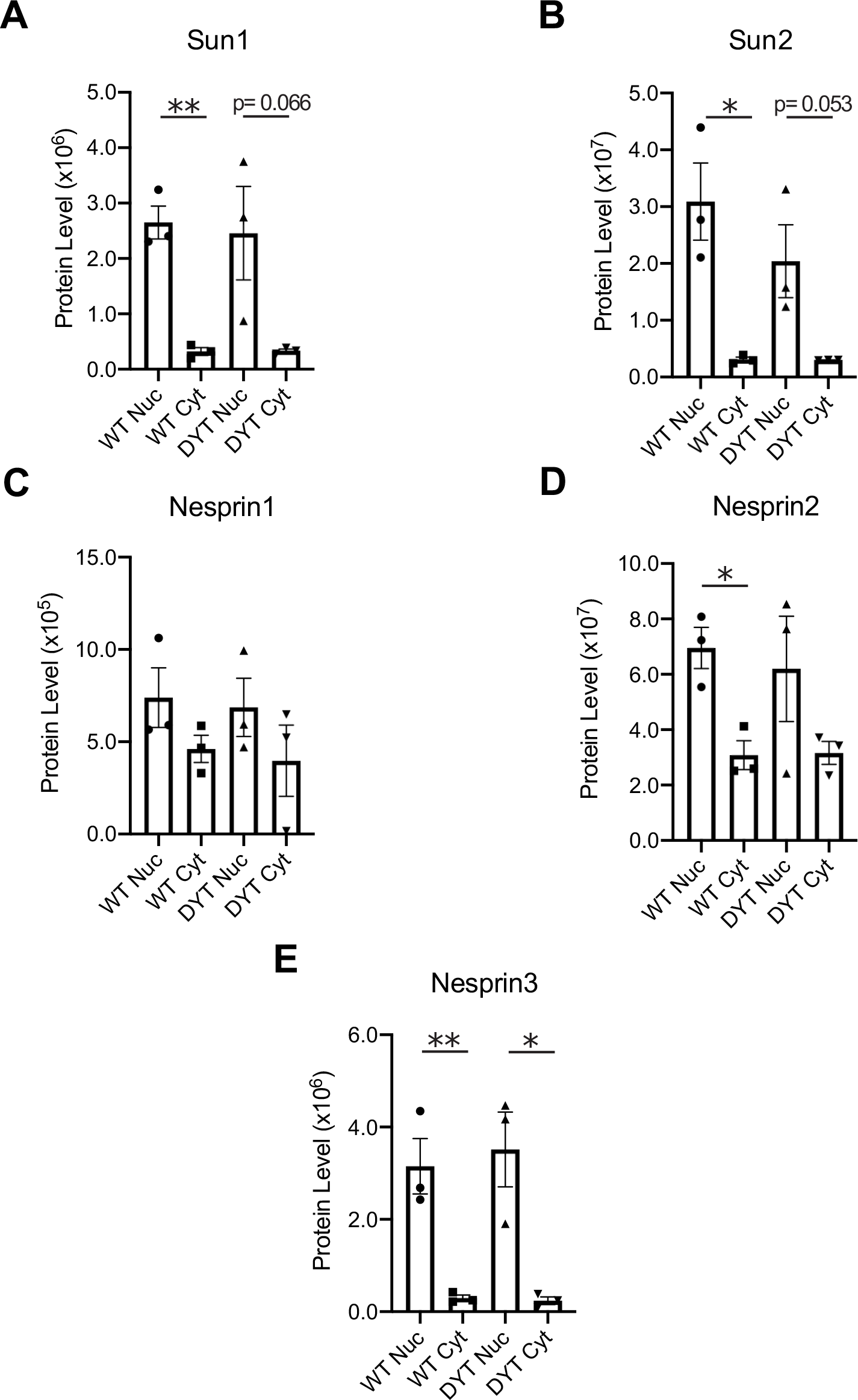
LINC proteins fractionation across genotype following Veh treatment. Protein levels in the nuclear and cytosolic fractions of both WT and DYT-TOR1A cell lines for (A) Sun1, (B) Sun2, (C) Nesprin1, (D) Nesprin2, and (E) Nesprin3. Significance was determined by unpaired t-test (n=3 biological replicates; *p<0.05; **p<0.01).

**Fig. S5.**
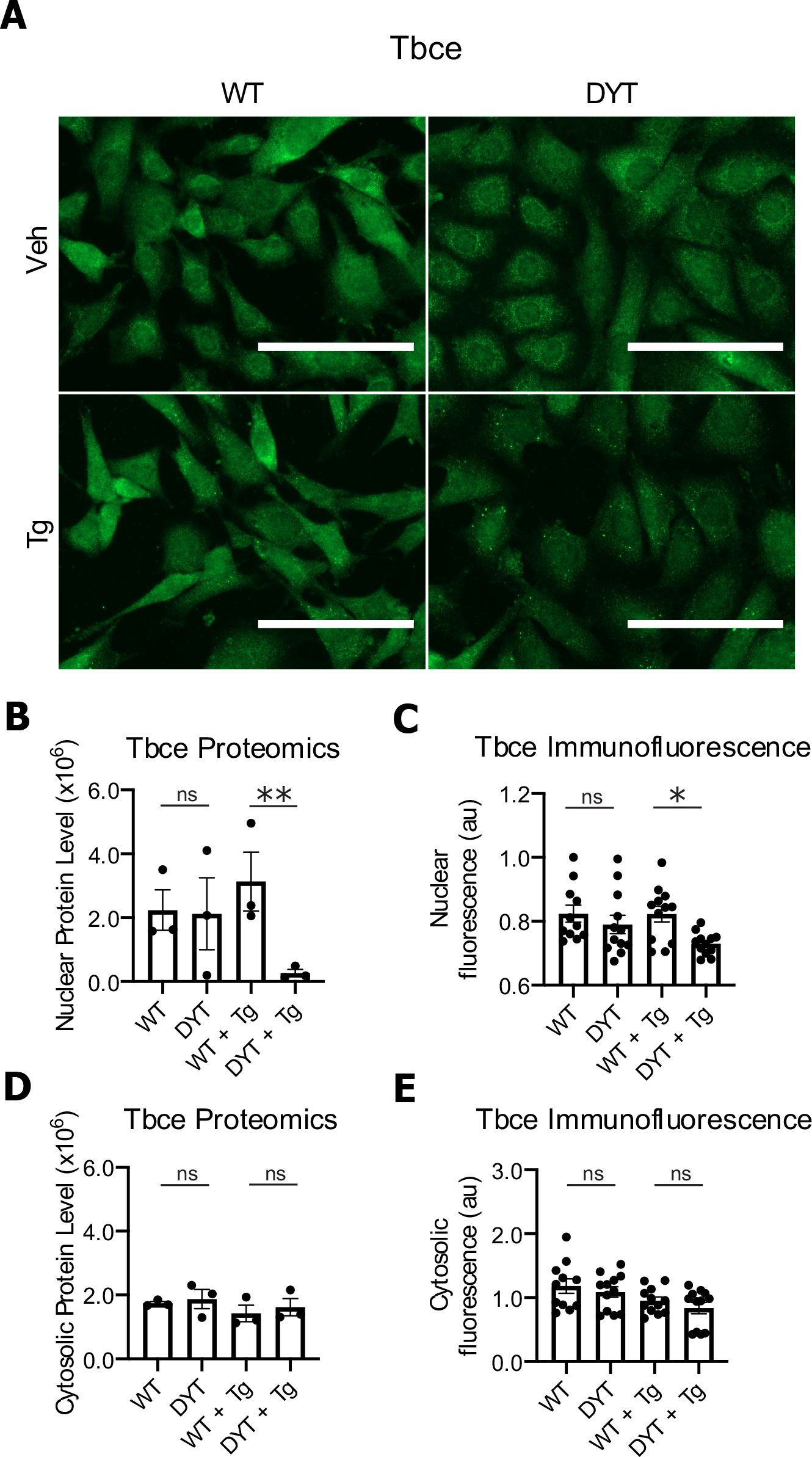
Quantification of Tbce protein by immunocytochemical staining. (A) Representative images of Tbce immunofluorescence in WT and DYT-TOR1A MEF lines treated with either Veh or Tg. (Scale bar = 100 µm) (B) Quantitative proteomics data on Tbce protein levels within the nuclear fractions. (C) Quantification of Tbce immunofluorescence within the nucleus. (D) Quantitative proteomics data on Tbce protein levels within the cytosolic fractions. (E) Quantification of Tbce immunofluorescence within the cytosol. For the proteomics data, significance was determined by unpaired t-test (n=3 biological replicates; **p<0.01). For the immunofluorescence data, significance was determined by unpaired t-test (n=12 biological replicates with 4 distinct wells being quantified for each of the three unique cell lines per genotype; *p<0.05).

**Fig. S6.**
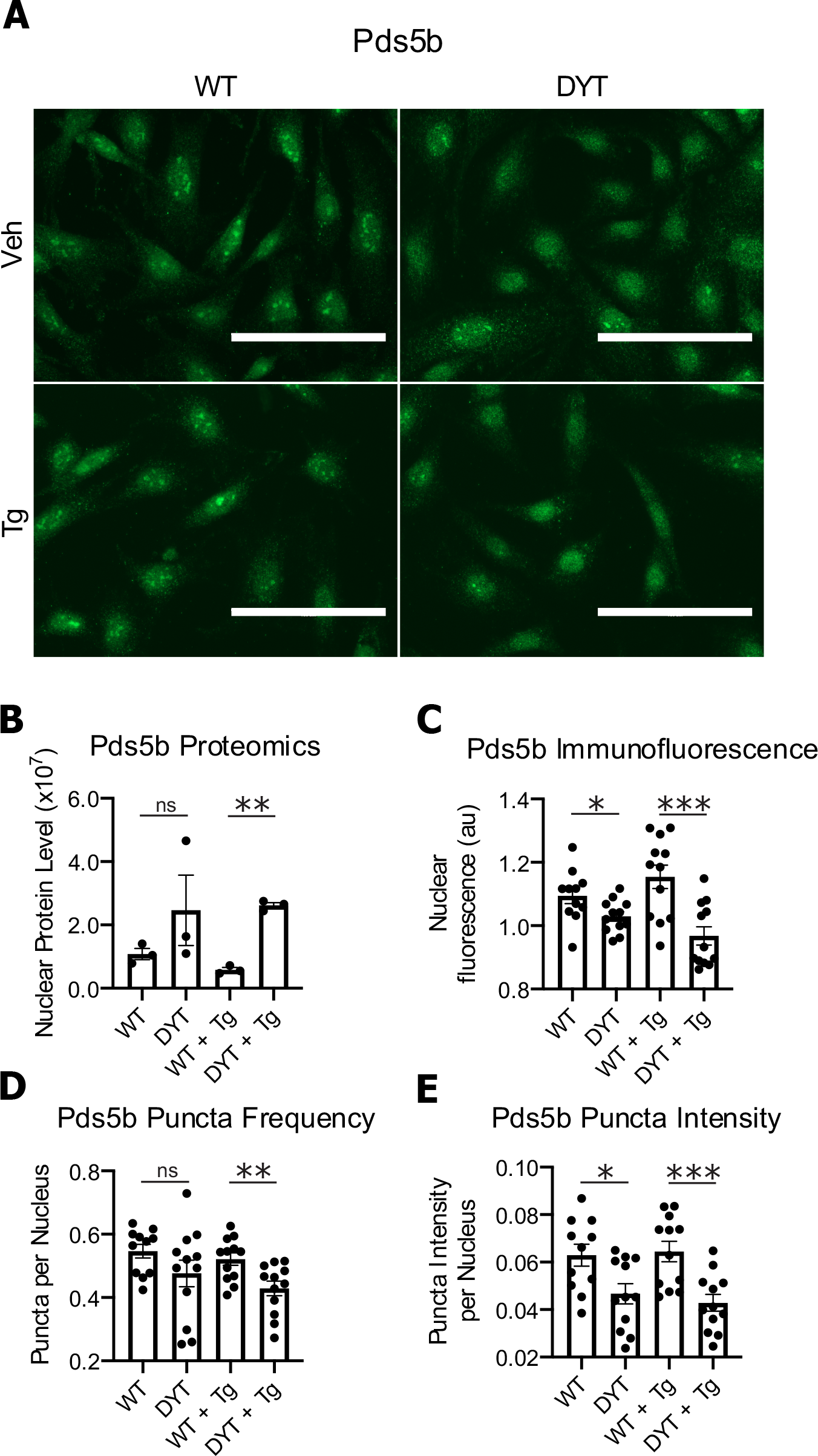
Quantification of Pds5b protein by immunocytochemical staining. (A) Representative images of Pds5b immunofluorescence in WT and DYT-TOR1A MEF lines treated with either Veh or Tg. (Scale bar = 100 µm) (B) Quantitative proteomics data on Pds5b protein levels within the nuclear fractions. (C) Quantification of Pds5b immunofluorescence within the nucleus. (D) Quantification of Pds5b puncta frequency within the nucleus. (E) Quantification of cumulative Pds5b puncta intensity within the cell nucleus. For the proteomics data, significance was determined by unpaired t-test (n=3 biological replicates; **p<0.01). For the immunofluorescence data, significance was determined by unpaired t-test (n=12 biological replicates with 4 distinct wells being quantified for each of the three unique cell lines per genotype; *p<0.05, ***p<0.001).

**Fig. S7.**
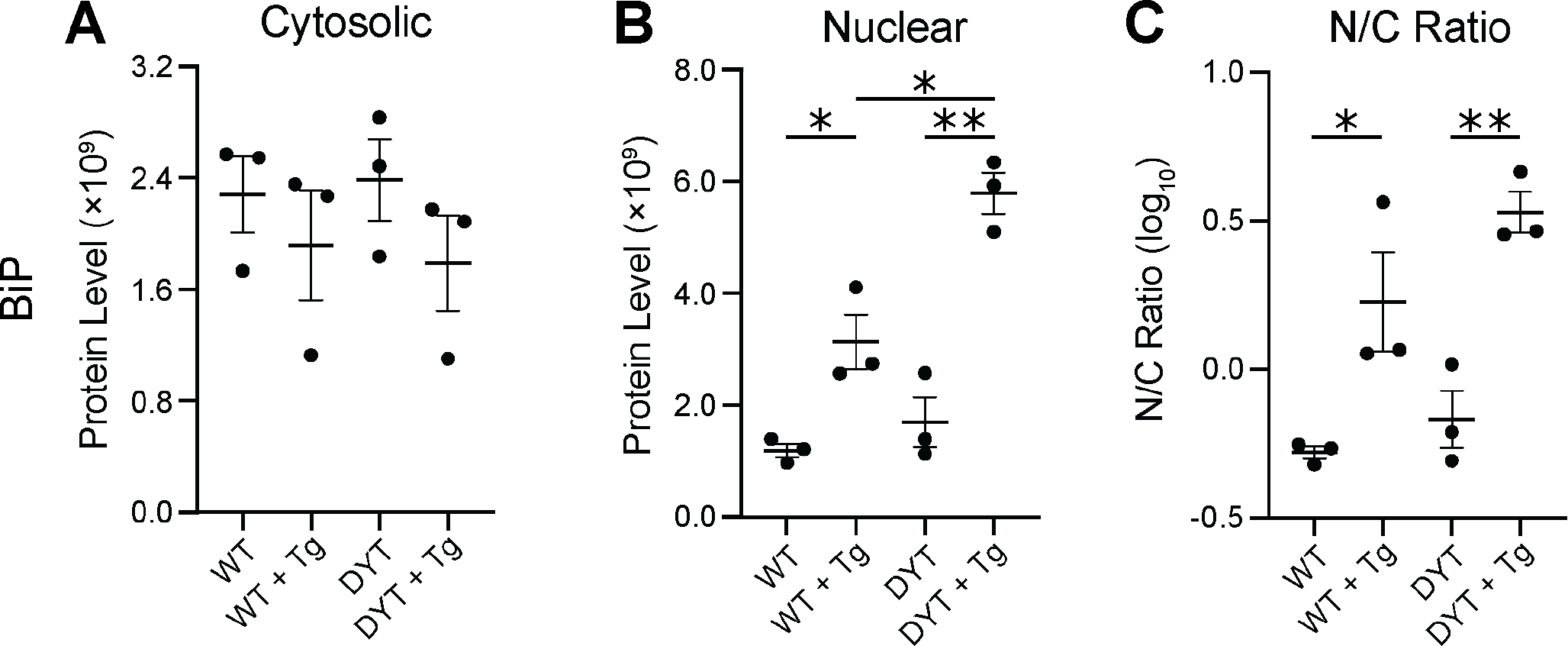
Genotype and Tg stress effects on subcellular fractionation of BiP. (A-B) LC/MS/MS quantified relative protein abundance of BiP in the cytosolic (A) and nuclear (B) fraction. (C) Ratio of nuclear/cytosolic protein levels. Significance was determined via unpaired t-test (n=3 biological replicates; *p<0.05; **p<0.01).

## References

Balint, B., Mencacci, N. E., Valente, E. M., Pisani, A., Rothwell, J., Jankovic, J., Vidailhet, M., & Bhatia, K. P. (2018). Dystonia. Nature Reviews Disease Primers, 4(1), 1–23. https://doi.org/10.1038/s41572-018-0023-6

Beauvais, G., Bode, N. M., Watson, J. L., Wen, H., Glenn, K. A., Kawano, H., Charles Harata, N., Ehrlich, M. E., & Gonzalez-Alegre, P. (2016). Disruption of Protein Processing in the Endoplasmic Reticulum of DYT1 Knock-in Mice Implicates Novel Pathways in Dystonia Pathogenesis. https://doi.org/10.1523/JNEUROSCI.0669-16.2016

Beauvais, G., Rodriguez-Losada, N., Ying, L., Zakirova, Z., Watson, J. L., Readhead, B., Gadue, P., French, D. L., Ehrlich, M. E., & Gonzalez-Alegre, P. (2018). Exploring the Interaction Between eIF2α Dysregulation, Acute Endoplasmic Reticulum Stress and DYT1 Dystonia in the Mammalian Brain. Neuroscience, 371, 455–468. https://doi.org/10.1016/j.neuroscience.2017.12.033

Bragg, D. C., Kaufman, C. A., Kock, N., & Breakefield, X. O. (2004). Inhibition of N-linked glycosylation prevents inclusion formation by the dystonia-related mutant form of torsinA. Molecular and Cellular Neuroscience, 27(4), 417–426. https://doi.org/10.1016/j.mcn.2004.07.009

Bressman, S. B. (2004). Dystonia genotypes, phenotypes, and classification. In Advances in neurology.

Calakos, N., Patel, V. D., Gottron, M., Wang, G., Tran-Viet, K. N., Brewington, D., Beyer, J. L., Steffens, D. C., Krishnan, R. R., & Züchner, S. (2010). Functional evidence implicating a novel TOR1A mutation in idiopathic, late-onset focal dystonia. Journal of Medical Genetics, 47(9), 646–650. https://doi.org/10.1136/jmg.2009.072082

Cao, S., Gelwix, C. C., Caldwell, K. A., & Caldwell, G. A. (2005). Torsin-mediated protection from cellular stress in the dopaminergic neurons of Caenorhabditis elegans. Journal of Neuroscience, 25(15), 3801–3812. https://doi.org/10.1523/JNEUROSCI.5157-04.2005

Carr, S., Aebersold, R., Baldwin, M., Burlingame, A., Clauser, K., & Nesvizhskii, A. (2004). The need for guidelines in publication of peptide and protein identification data: Working group on publication guidelines for peptide and protein identification data. In Molecular and Cellular Proteomics (Vol. 3, Issue 6, pp. 531–532). American Society for Biochemistry and Molecular Biology. https://doi.org/10.1074/mcp.T400006-MCP200

Chalfant, M., Barber, K. W., Borah, S., Thaller, D., & Lusk, C. P. (2019). Expression of TorsinA in a heterologous yeast system reveals interactions with lumenal domains of LINC and nuclear pore complex components. Molecular Biology of the Cell, 30(5), 530–541. https://doi.org/10.1091/mbc.E18-09-0585

Chase, A. R., Laudermilch, E., Wang, J., Shigematsu, H., Yokoyama, T., & Schlieker, C. (2017). Dynamic functional assembly of the Torsin AAA+ ATPase and its modulation by LAP1. Molecular Biology of the Cell. https://doi.org/10.1091/mbc.E17-05-0281

Chen, P., Burdette, A. J., Porter, J. C., Ricketts, J. C., Fox, S. A., Nery, F. C., Hewett, J. W., Berkowitz, L. A., Breakefield, X. O., Caldwell, K. A., & Caldwell, G. A. (2010). The early-onset torsion dystonia-associated protein, torsinA, is a homeostatic regulator of endoplasmic reticulum stress response. Human Molecular Genetics, 19(18), 3502–3515. https://doi.org/10.1093/hmg/ddq266

Cho, J. A., Zhang, X., Miller, G. M., Lencer, W. I., & Nery, F. C. (2014). 4-Phenylbutyrate Attenuates the ER Stress Response and Cyclic AMP Accumulation in DYT1 Dystonia Cell Models. PLoS ONE, 9(11), e110086. https://doi.org/10.1371/journal.pone.0110086

Ding, B., Tang, Y., Ma, S., Akter, M., Liu, M. L., Zang, T., & Zhang, C. L. (2021). Disease modeling with human neurons reveals lmnb1 dysregulation underlying dyt1 dystonia. Journal of Neuroscience. https://doi.org/10.1523/JNEUROSCI.2507-20.2020

Enenkel, C. (2014). Nuclear transport of yeast proteasomes. In Biomolecules (Vol. 4, Issue 4, pp. 940–955). MDPI AG. https://doi.org/10.3390/biom4040940

Esra Demircioglu, F., Sosa, B. A., Ingram, J., Ploegh, H. L., & Schwartz, T. U. (2016). Structures of torsinA and its disease-mutant complexed with an activator reveal the molecular basis for primary dystonia. ELife. https://doi.org/10.7554/eLife.17983

Gonzalez-Alegre, P., & Paulson, H. L. (2004). Aberrant Cellular Behavior of Mutant TorsinA Implicates Nuclear Envelope Dysfunction in DYT1 Dystonia. Journal of Neuroscience, 24(11), 2593–2601. https://doi.org/10.1523/JNEUROSCI.4461-03.2004

Goodchild, R. E., Buchwalter, A. L., Naismith, T. V., Holbrook, K., Billion, K., Dauer, W. T., Liang, C. C., Dear, M. L., & Hanson, P. I. (2015). Access of torsinA to the inner nuclear membrane is activity dependent and regulated in the endoplasmic reticulum. Journal of Cell Science, 128(15), 2854–2865. https://doi.org/10.1242/jcs.167452

Goodchild, R. E., & Dauer, W. T. (2004). Mislocalization to the nuclear envelope: An effect of the dystonia-causing torsinA mutation. Proceedings of the National Academy of Sciences of the United States of America, 101(3), 847–852. https://doi.org/10.1073/pnas.0304375101

Goodchild, R. E., & Dauer, W. T. (2005). The AAA+ protein torsinA interacts with a conserved domain present in LAP1 and a novel ER protein. Journal of Cell Biology, 168(6), 855–862. https://doi.org/10.1083/jcb.200411026

Goodchild, R. E., Kim, C. E., & Dauer, W. T. (2005). Loss of the dystonia-associated protein torsinA selectively disrupts the neuronal nuclear envelope. Neuron, 48(6), 923–932. https://doi.org/10.1016/j.neuron.2005.11.010

György, B., Cruz, L., Yellen, D., Aufiero, M., Alland, I., Zhang, X., Ericsson, M., Fraefel, C., Li, C., Takeda, S., Hyman, B. T., & Breakefield, X. O. (2018). Mutant torsinA in the heterozygous DYT1 state compromises HSV propagation in infected neurons and fibroblasts. Scientific Reports, 8(1). https://doi.org/10.1038/s41598-018-19865-2

Harding, H. (2003). Immortalization of MEF with SV40 T antigen. In Internet.

Harding, H. P., Zhang, Y., Bertolotti, A., Zeng, H., & Ron, D. (2000). Perk is essential for translational regulation and cell survival during the unfolded protein response. Molecular Cell, 5(5), 897–904. https://doi.org/10.1016/S1097-2765(00)80330-5

Hewett, J., Gonzalez-Agosti, C., Slater, D., Ziefer, P., Li, S., Bergeron, D., Jacoby, D. J., Ozelius, L. J., Ramesh, V., & Breakefield, X. O. (2000). Mutant torsinA, responsible for early-onset torsion dystonia, forms membrane inclusions in cultured neural cells. Human Molecular Genetics. https://doi.org/10.1093/hmg/9.9.1403

Jankovic, J., & Tintner, R. (2001). Dystonia and parkinsonism. Parkinsonism and Related Disorders, 8(2), 109–121. https://doi.org/10.1016/S1353-8020(01)00025-6

Jokhi, V., Ashley, J., Nunnari, J., Noma, A., Ito, N., Wakabayashi-Ito, N., Moore, M. J., & Budnik, V. (2013). Torsin Mediates Primary Envelopment of Large Ribonucleoprotein Granules at the Nuclear Envelope. Cell Reports. https://doi.org/10.1016/j.celrep.2013.03.015

Jozefczuk, J., Drews, K., & Adjaye, J. (2012). Preparation of mouse embryonic fibroblast cells suitable for culturing human embryonic and induced pluripotent stem cells. Journal of Visualized Experiments. https://doi.org/10.3791/3854

Jungwirth, M., Dear, M. L., Brown, P., Holbrook, K., & Goodchild, R. (2010). Relative tissue expression of homologous torsinB correlates with the neuronal specific importance of DYT1 dystonia-associated torsinA. Human Molecular Genetics, 19(5), 888–900. https://doi.org/10.1093/hmg/ddp557

Jungwirth, M. T., Kumar, D., Jeong, D. Y., & Goodchild, R. E. (2011). The nuclear envelope localization of DYT1 dystonia torsinA-ΔE requires the SUN1 LINC complex component. BMC Cell Biology, 12. https://doi.org/10.1186/1471-2121-12-24

Khalil, B., Morderer, D., Price, P. L., Liu, F., & Rossoll, W. (2018). mRNP assembly, axonal transport, and local translation in neurodegenerative diseases. In Brain Research (Vol. 1693, Issue Pt A, pp. 75–91). Elsevier B.V. https://doi.org/10.1016/j.brainres.2018.02.018

Kim, C. E., Perez, A., Perkins, G., Ellisman, M. H., & Dauer, W. T. (2010). A molecular mechanism underlying the neural-specific defect in torsinA mutant mice. Proceedings of the National Academy of Sciences of the United States of America. https://doi.org/10.1073/pnas.0912877107

Kim, J. E., Hong, Y. H., Kim, J. Y., Jeon, G. S., Jung, J. H., Yoon, B. N., Son, S. Y., Lee, K. W., Kim, J. Il, & Sung, J. J. (2017). Altered nucleocytoplasmic proteome and transcriptome distributions in an in vitro model of amyotrophic lateral sclerosis. PLoS ONE, 12(4). https://doi.org/10.1371/journal.pone.0176462

Kustedjo, K., Bracey, M. H., & Cravatt, B. F. (2000). Torsin A and its Torsion Dystonia-Associated Mutant Form Are Lumenal Glycoproteins that Exhibit Distinct Subcellular Localizations Running title: Biochemical characterization of torsin A. JBC Papers in Press.

Laudermilch, E., Tsai, P. L., Graham, M., Turner, E., Zhao, C., & Schlieker, C. (2016). Dissecting Torsin/cofactor function at the nuclear envelope: A genetic study. Molecular Biology of the Cell, 27(25), 3964–3971. https://doi.org/10.1091/mbc.E16-07-0511

Li, J., Liang, C. C., Pappas, S. S., & Dauer, W. T. (2020). TorsinB overexpression prevents abnormal twisting in DYT1 dystonia mouse models. ELife, 9. https://doi.org/10.7554/eLife.54285

Liang, C. C., Tanabe, L. M., Jou, S., Chi, F., & Dauer, W. T. (2014). TorsinA hypofunction causes abnormal twisting movements and sensorimotor circuit neurodegeneration. Journal of Clinical Investigation, 124(7), 3080–3092. https://doi.org/10.1172/JCI72830

Martin, J. N., Bair, T. B., Bode, N., Dauer, W. T., & Gonzalez-Alegre, P. (2009). Transcriptional and proteomic profiling in a cellular model of DYT1 dystonia. Neuroscience, 164(2), 563– 572. https://doi.org/10.1016/j.neuroscience.2009.07.068

McQuin, C., Goodman, A., Chernyshev, V., Kamentsky, L., Cimini, B. A., Karhohs, K. W., Doan, M., Ding, L., Rafelski, S. M., Thirstrup, D., Wiegraebe, W., Singh, S., Becker, T., Caicedo, J. C., & Carpenter, A. E. (2018). CellProfiler 3.0: Next-generation image processing for biology. PLoS Biology. https://doi.org/10.1371/journal.pbio.2005970

Mootha, V. K., Lindgren, C. M., Eriksson, K. F., Subramanian, A., Sihag, S., Lehar, J., Puigserver, P., Carlsson, E., Ridderstråle, M., Laurila, E., Houstis, N., Daly, M. J., Patterson, N., Mesirov, J. P., Golub, T. R., Tamayo, P., Spiegelman, B., Lander, E. S., Hirschhorn, J. N., … Groop, L. C. (2003). PGC-1α-responsive genes involved in oxidative phosphorylation are coordinately downregulated in human diabetes. Nature Genetics. https://doi.org/10.1038/ng1180

Naismith, T. V., Heuser, J. E., Breakefield, X. O., & Hanson, P. I. (2004). TorsinA in the nuclear envelope. Proceedings of the National Academy of Sciences of the United States of America, 101(20), 7612–7617. https://doi.org/10.1073/pnas.0308760101

Nery, F. C., Armata, I. A., Farley, J. E., Cho, J. A., Yaqub, U., Chen, P., Da Hora, C. C., Wang, Q., Tagaya, M., Klein, C., Tannous, B., Caldwell, K. A., Caldwell, G. A., Lencer, W. I., Ye, Y., & Breakefield, X. O. (2011). TorsinA participates in endoplasmic reticulum-associated degradation. Nature Communications, 2(1). https://doi.org/10.1038/ncomms1383

Nery, F. C., Zeng, J., Niland, B. P., Hewett, J., Farley, J., Irimia, D., Li, Y., Wiche, G., Sonnenberg, A., & Breakefield, X. O. (2008). TorsinA binds the KASH domain of nesprins and participates in linkage between nuclear envelope and cytoskeleton. Journal of Cell Science, 121(20), 3476–3486. https://doi.org/10.1242/jcs.029454

Ortega, J. A., Daley, E. L., Kour, S., Samani, M., Tellez, L., Smith, H. S., Hall, E. A., Esengul, Y. T., Tsai, Y. H., Gendron, T. F., Donnelly, C. J., Siddique, T., Savas, J. N., Pandey, U. B., & Kiskinis, E. (2020). Nucleocytoplasmic Proteomic Analysis Uncovers eRF1 and Nonsense-Mediated Decay as Modifiers of ALS/FTD C9orf72 Toxicity. Neuron, 106(1), 90–107.e13. https://doi.org/10.1016/j.neuron.2020.01.020

Ozelius, L. J., Hewett, J. W., Page, C. E., Bressman, S. B., Kramer, P. L., Shalish, C., De Leon, D., Brin, M. F., Raymond, D., Corey, D. P., Fahn, S., Risch, N. J., Buckler, A. J., Gusella, J. F., & Breakefield, X. O. (1997). The early-onset torsion dystonia gene (DYT1) encodes an ATP-binding protein. Nature Genetics. https://doi.org/10.1038/ng0997-40

Pappas, S. S., Liang, C. C., Kim, S., Rivera, C. A. O., & Dauer, W. T. (2018). TorsinA dysfunction causes persistent neuronal nuclear pore defects. Human Molecular Genetics. https://doi.org/10.1093/hmg/ddx405

Rampello, A. J., Laudermilch, E., Vishnoi, N., Prohet, S. M., Zhao, C., Patrick Lusk, C., & Schlieker, C. (2019). Torsin ATPases are required to complete nuclear pore complex biogenesis in interphase 1 *2*. https://doi.org/10.1101/821835

Rittiner, J. E., Caffall, Z. F., Hernández-Martinez, R., Sanderson, S. M., Pearson, J. L., Tsukayama, K. K., Liu, A. Y., Xiao, C., Tracy, S., Shipman, M. K., Hickey, P., Johnson, J., Scott, B., Stacy, M., Saunders-Pullman, R., Bressman, S., Simonyan, K., Sharma, N., Ozelius, L. J., … Calakos, N. (2016). Functional Genomic Analyses of Mendelian and Sporadic Disease Identify Impaired eIF2α Signaling as a Generalizable Mechanism for Dystonia. Neuron, 92(6), 1238–1251. https://doi.org/10.1016/j.neuron.2016.11.012

Ron, D. (2002). Translational control in the endoplasmic reticulum stress response. Journal of Clinical Investigation, 110(10), 1383–1388. https://doi.org/10.1172/jci16784

Ross Buchan, J. (2014). MRNP granules Assembly, function, and connections with disease. In RNA Biology (Vol. 11, Issue 8, pp. 1019–1030). Landes Bioscience. https://doi.org/10.4161/15476286.2014.972208

Saunders, C. A., Harris, N. J., Willey, P. T., Woolums, B. M., Wang, Y., McQuown, A. J., Schoenhofen, A., Worman, H. J., Dauer, W. T., Gundersen, G. G., & Luxton, G. W. G. (2017). TorsinA controls TAN line assembly and the retrograde flow of dorsal perinuclear actin cables during rearward nuclear movement. The Journal of Cell Biology, 216(3), 657– 674. https://doi.org/10.1083/jcb.201507113

Speese, S. D., Ashley, J., Jokhi, V., Nunnari, J., Barria, R., Li, Y., Ataman, B., Koon, A., Chang, Y. T., Li, Q., Moore, M. J., & Budnik, V. (2012). Nuclear envelope budding enables large ribonucleoprotein particle export during synaptic Wnt signaling. Cell, 149(4), 832–846. https://doi.org/10.1016/j.cell.2012.03.032

Subramanian, A., Tamayo, P., Mootha, V. K., Mukherjee, S., Ebert, B. L., Gillette, M. A., Paulovich, A., Pomeroy, S. L., Golub, T. R., Lander, E. S., & Mesirov, J. P. (2005). Gene set enrichment analysis: A knowledge-based approach for interpreting genome-wide expression profiles. Proceedings of the National Academy of Sciences of the United States of America, 102(43), 15545–15550. https://doi.org/10.1073/pnas.0506580102

Suzuki, K., Bose, P., Leong-Quong, R. Y., Fujita, D. J., & Riabowol, K. (2010). REAP: A two minute cell fractionation method. BMC Research Notes. https://doi.org/10.1186/1756-0500-3-294

Tanabe, L. M., Liang, C. C., & Dauer, W. T. (2016). Neuronal Nuclear Membrane Budding Occurs during a Developmental Window Modulated by Torsin Paralogs. Cell Reports, 16(12), 3322–3333. https://doi.org/10.1016/j.celrep.2016.08.044

Tarsy, D., & Simon, D. K. (2006). Dystonia. New England Journal of Medicine. https://doi.org/10.1056/NEJMra055549

Torres, G. E., Sweeney, A. L., Beaulieu, J. M., Shashidharan, P., & Caron, M. G. (2004). Effect of torsinA on membrane proteins reveals a loss of function and a dominant-negative phenotype of the dystonia-associated ΔE-torsinA mutant. Proceedings of the National Academy of Sciences of the United States of America, 101(44), 15650–15655. https://doi.org/10.1073/pnas.0308088101

Tribl, F., Gerlach, M., Marcus, K., Asan, E., Tatschner, T., Arzberger, T., Meyer, H. E., Bringmann, G., & Riederer, P. (2005). “Subcellular proteomics” of neuromelanin granules isolated from the human brain. In Molecular and Cellular Proteomics (Vol. 4, Issue 7, pp. 945–957). American Society for Biochemistry and Molecular Biology Inc. https://doi.org/10.1074/mcp.M400117-MCP200

van Harten, P. N., Hoek, H. W., & Kahn, R. S. (1999). Fortnightly review: Acute dystonia induced by drug treatment. BMJ, 319(7210), 623–626. https://doi.org/10.1136/bmj.319.7210.623

Vander Heyden, A. B., Naismith, T. V., Snapp, E. L., Hodzic, D., & Hanson, P. I. (2009). LULL1 retargets torsinA to the nuclear envelope revealing an activity that is impaired by the DYT1 dystonia mutation. Molecular Biology of the Cell, 20(11), 2661–2672. https://doi.org/10.1091/mbc.E09-01-0094

VanGompel, M. J. W., Nguyen, K. C. Q., Hall, D. H., Dauer, W. T., & Rose, L. S. (2015). A novel function for the Caenorhabditis elegans torsin OOC-5 in nucleoporin localization and nuclear import. Molecular Biology of the Cell. https://doi.org/10.1091/mbc.E14-07-1239

Vasudevan, A., Breakefield, X. O., & Bhide, P. G. (2006). Developmental patterns of torsinA and torsinB expression. Brain Research, 1073–1074(1), 139–145. https://doi.org/10.1016/j.brainres.2005.12.087

Wühr, M., Güttler, T., Peshkin, L., McAlister, G. C., Sonnett, M., Ishihara, K., Groen, A. C., Presler, M., Erickson, B. K., Mitchison, T. J., Kirschner, M. W., & Gygi, S. P. (2015). The Nuclear Proteome of a Vertebrate. Current Biology, 25(20), 2663–2671. https://doi.org/10.1016/j.cub.2015.08.047

Zacchi, L. F., Wu, H. C., Bell, S. L., Millen, L., Paton, A. W., Paton, J. C., Thomas, P. J., Zolkiewski, M., & Brodsky, J. L. (2014). The bip molecular chaperone plays multiple roles during the biogenesis of torsina, an aaa atpase associated with the neurological disease early-onset torsion dystonia. Journal of Biological Chemistry, 289(18), 12727–12747. https://doi.org/10.1074/jbc.M113.529123

Zakirova, Z., Fanutza, T., Bonet, J., Readhead, B., Zhang, W., Yi, Z., Beauvais, G., Zwaka, T. P., Ozelius, L. J., Blitzer, R. D., Gonzalez-Alegre, P., & Ehrlich, M. E. (2018). Mutations in THAP1/DYT6 reveal that diverse dystonia genes disrupt similar neuronal pathways and functions. PLoS Genetics, 14(1). https://doi.org/10.1371/journal.pgen.1007169

Zhao, C., Brown, R. S. H., Chase, A. R., Eisele, M. R., & Schlieker, C. (2013). Regulation of Torsin ATPases by LAP1 and LULL1. Proceedings of the National Academy of Sciences of the United States of America, 110(17). https://doi.org/10.1073/pnas.1300676110

Zhao, C., Brown, R. S. H., Tang, C. H. A., Hu, C. C. A., & Schlieker, C. (2016). Site-specific proteolysis mobilizes TorsinA from the membrane of the endoplasmic reticulum (ER) in response to ER stress and B cell stimulation. Journal of Biological Chemistry, 291(18), 9469–9481. https://doi.org/10.1074/jbc.M115.709337

Zhou, Y., Zhou, B., Pache, L., Chang, M., Khodabakhshi, A. H., Tanaseichuk, O., Benner, C., & Chanda, S. K. (2019). Metascape provides a biologist-oriented resource for the analysis of systems-level datasets. Nature Communications. https://doi.org/10.1038/s41467-019-09234-6

